# Enhanced Visualization of Influenza A Virus Entry Using Virus-View Atomic Force Microscopy

**DOI:** 10.1101/2024.07.19.603848

**Authors:** Aiko Yoshida, Yoshitsugu Uekusa, Takeshi Suzuki, Michael Bauer, Nobuaki Sakai, Yohei Yamauchi

**Author notes:** These authors contributed equally to this work (Aiko Yoshida, Nobuaki Sakai). Novo Nordisk, Måløv, 2760, Denmark (Michael Bauer).

## Abstract

Virus entry begins with attachment of virions to the cell surface, multivalent binding of viral proteins to receptors, signaling, and endocytosis. Using ViViD-AFM (Virus-View Dual confocal and Atomic Force Microscopy), we visualized the nanoscale morphology of influenza A virus (IAV) virions interacting with the cell membrane during virus entry. Following attachment to the cell surface, spherical IAV (90-100 nm in diameter) diffused in a sialic acid- and neuraminidase-dependent manner. Reduced diffusion signified the onset of clathrin coat assembly, followed by formation of actin-rich ruffles that promoted pit closure and IAV endocytosis. Cell surface ruffles sheared filamentous IAV (>1µm in length) into shorter fragments that became internalized. ViViD-AFM is a powerful tool that provides nanoscale morphological insights of virus-cell membrane interplay in living cells.

---

The entry of a virus into a host cell begins with the initial encounter between the virus particle and the host cell surface. After attachment, the virus undergoes lateral diffusion, followed by internalization into vesicles via membrane invaginations (1, 2). Interactions between the viral receptor-binding surface proteins and cellular receptors are generally weak but multivalent, leading to high avidity. This multivalent binding results in receptor clustering, signaling, and the internalization of the virus particle (1, 2).

Influenza A virus (IAV) is an enveloped RNA virus approximately 100 nm in size. The viral envelope glycoprotein hemagglutinin (HA) binds to sialic acid receptors, which are abundant on the cell surface. Viral neuraminidase catalyzes the destruction of sialic acid receptors, regulating the attachment and detachment of IAV through these opposing activities (3, 4). Functional entry receptors for IAV include the epidermal growth factor receptor (EGFR), L-type voltage-dependent calcium channel alpha 1C subunit (Cav1.2), major histocompatibility complex (MHC) class II, and transferrin receptor (TfR) (5–8). IAV internalization is facilitated by clathrin-mediated endocytosis (CME), macropinocytosis, and dynamin-independent endocytic processes in non-coated invaginations (9–12).

The nanometer resolution of atomic force microscopy (AFM) is valuable for detailed morphological analysis of purified protein complexes and viral particles (13–16), as well as for time-lapse imaging of plasma membrane morphology, submembrane actin dynamics, and CME (17–21). In this study, we established a hybrid AFM system called ViViD-AFM (Virus-view Dual confocal and Atomic Force Microscopy), which employs a minimally invasive ultra-narrow cantilever with dimensions of 9.0 μm in length, 0.8 μm in width, and 0.1 μm in thickness (**Fig. 1A, S1**). By minimizing the cantilever’s width, we reduced the cantilever peak force while maintaining scanning speed. This unique design reduced the cantilever spring constant to 0.04 N/m, low enough to preserve delicate virus-receptor interactions. Consequently, ViViD-AFM has a field of view of 27 µm², enabling the analysis of IAV cell attachment, diffusion, endocytosis, and cell membrane dynamics at nanometer resolution in living cells with a time resolution of 5 s.

**Fig. 1.**
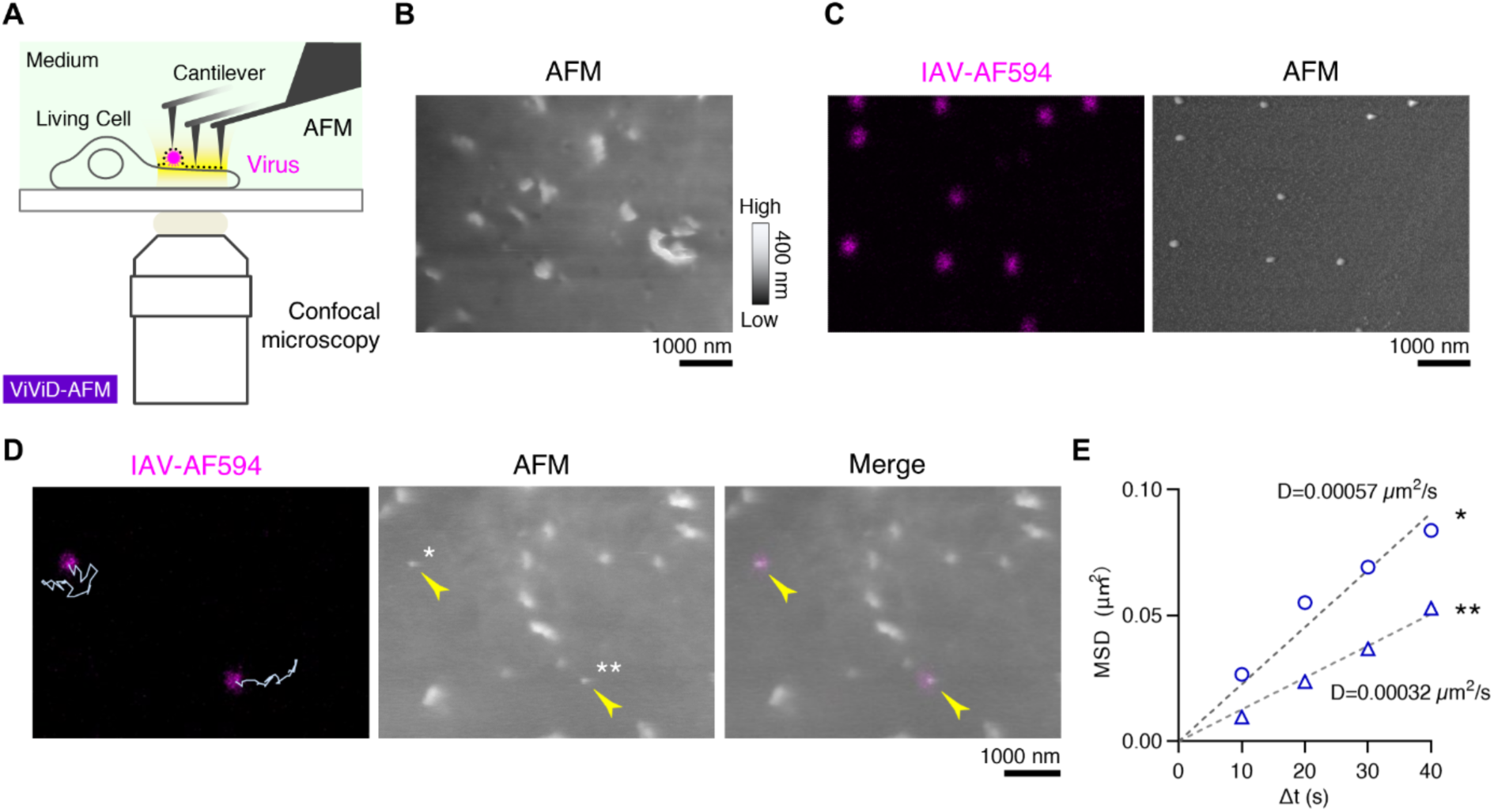
Establishment of ViViD-AFM for IAV cell entry studies in MDCK cells. **(A)** Schematic of ViViD-AFM for simultaneous dual live imaging of morphology and fluorescence. The cantilever scans the surface of the cell and the IAV virion, providing morphology images, while the confocal microscope detects the fluorescence. **(B)** Morphology image of living cell surface acquired by AFM. The AFM image is a topographic image, with the height direction: bright areas are higher and dark areas are lower. **(C)** Fluorescence (left) and morphology image (right) of glass-bound IAV virions labeled with Alexa Fluor 594 (IAV-AF594) acquired by ViViD-AFM. **(D)** Time-lapse ViViD-AFM imaging of cell surface IAV virions (arrowheads) at 27°C at 10 s intervals. Fluorescence (left), morphology (middle), and merged images (right) are shown. Two IAV virions were tracked and their diffusion trajectories over 300 s were superimposed in the image. **(E)** Quantification of IAV diffusion. Mean square displacement (MSD) plot of each of the two virions (* and **) in panel (D) was overlaid with dashed lines indicating two-dimensional free diffusion. Diffusion coefficient was calculated from MSD plots over 30 frames (300 s). D, diffusion coefficient. (B–D) Scale bars: 1000 nm.

## Establishment of ViViD-AFM for IAV cell entry studies in MDCK cells

To evaluate if ViViD-AFM was suitable for detecting the cell surface morphology of live cultured cells, we used Madin-Darby canine kidney (MDCK) cells which are widely used for IAV isolation and vaccine production due to their high susceptibility to infection by various IAV strains (22). MDCK cells stably expressing the filamentous actin marker EGFP-Lifeact were imaged using ViViD-AFM. The cell surface morphology within the field of view (6.5 µm × 4.0 µm) revealed numerous membrane protrusions (**Fig. 1B, Movie S1**). ViViD-AFM imaging showed that the membrane protrusions were 100-200 nm in height and correlated with EGFP-Lifeact fluorescence intensity (r=0.7185) and hence were actin-rich membrane ruffles (**Fig. S2**). We next imaged glass-bound Alexa Fluor 594-labeled influenza A WSN (H1N1) (IAV-AF594) virions using ViViD-AFM (**Fig. 1C**). Analysis of 126 virions by ViViD-AFM showed 100 % fluorescent labeling efficiency and a mean virion height of 93.3 ± 7.5 nm (**Fig. S3**). These experiments established that ViViD-AFM was suitable for nanoscale imaging of living tissue culture cells and virus particles.

We next set out to establish ViViD-AFM for IAV-AF594 cell entry studies in live MDCK cells. At 27°C the binding forces between the viral HA and its sialic acid receptor is approximately 10-25 pN as measured by single-virus force spectroscopy (23). We evaluated if the ultra-narrow cantilever (**Fig. S1**) did not disrupt such delicate virus-receptor interactions. MDCK cells were inoculated with IAV-AF594 at a multiplicity of infection (MOI) of 100 and subjected to ViViD-AFM 5 min later. As shown in Fig. S4, we captured the IAV-AF594 fluorescent signal and morphology simultaneously over multiple time points, from the process of IAV attachment to subsequent lateral diffusion on the cell surface. To rule out effects of the cantilever on IAV diffusion, IAV-AF594-inoculated MDCK cells were first imaged with confocal microscopy alone (Step I, **Fig. S5A**), followed by dual imaging using ViViD-AFM (Step II, **Fig. S5A**). We compared the cell surface diffusion coefficients of 14 individual IAV particles and observed no difference between Steps I and II (**Fig. S5B**). This observation showed that ViViD-AFM was suitable for detecting physiological diffusion of virions on the cell surface. We next inoculated MDCK cells with IAV-AF594 at a MOI of 100 followed by ViViD-AFM imaging (**Movie S2**). The trajectory of two individual IAV-AF594 particles was analyzed by single particle tracking (**Fig. 1D**), and the diffusion coefficients (D) calculated from the mean squared displacement (MSD) plots for each virion derived were 0.00057 and 0.00032 µm^2^/s, respectively (**Fig. 1E**).

## IAV diffusion depends on viral neuraminidase and sialic acids

Lateral IAV diffusion is influenced by multivalent interactions between viral HA and sialic acid receptors (23–27), and the viral diffusion coefficient depends on the density of the sialic acids on the cell surface (24, 28). In addition, neuraminidase activity has been reported to influence IAV virion diffusion on the cell surface. First, we investigated how a single IAV virion diffuses over time. MDCK cells were inoculated with IAV-AF594 at an MOI of 100 at 37°C, followed by ViViD-AFM imaging at 5 s intervals. A single IAV virion that underwent diffusion for 1500 s on the cell surface was tracked (**Fig. 2A, Movie S3**). The 150 s simple moving average of the IAV diffusion log_10_D (the common logarithm of diffusion coefficient D) was −2.69 ± 0.23 log_10_ μm^2^/s (**Fig. 2B**). IAV diffusion coefficients showed microscale fluctuations depending on the region of the cell surface (**Fig. 2B, C**).

**Fig. 2.**
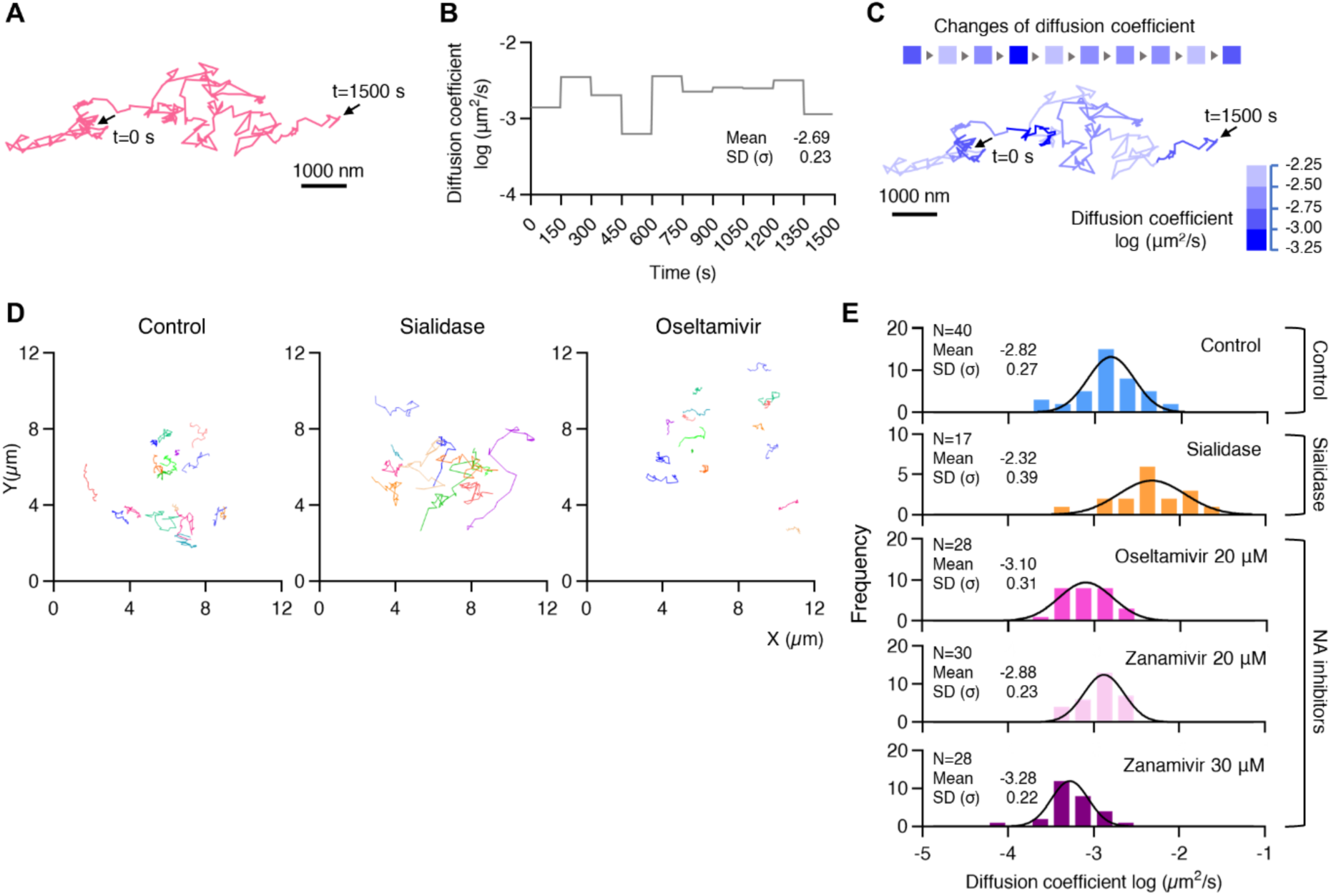
Sialic acids and neuraminidase regulate IAV cell surface diffusion. **(A)** Trajectory of a single IAV virion under Control condition from time-lapse ViViD-AFM imaging at 37°C over 1500 s at 5 s intervals. **(B)** Plot of diffusion coefficient of a single IAV virion in panel (A) over time. **(C)** Microscale-fluctuation of diffusion coefficient of an IAV virion: changes in cell surface diffusion coefficient of the virion over time (top) and corresponding trajectory (bottom) represented with color scale for diffusion coefficients. **(D)** Representative trajectories of IAV virions from multiple cells under Control (left), Sialidase (middle), and 20 µM Oseltamivir (right)-treated conditions acquired from time-lapse ViViD-AFM imaging at 37°C over 150 s at 5 s intervals. Each virion’s trajectory is colored differently. **(E)** Diffusion coefficient histograms of the of IAV virions on the cell surface. Top to bottom: Control (N=40), Sialidase (=17), 20 µM Oseltamivir (N=28), and Zanamivir at 20 µM (N=30) and 30 µM (N=28). Each histogram fitted with a normal distribution. (A, C) Scale bars: 1000 nm. (B, C, E) Diffusion coefficient was calculated from MSD plots over 30 frames (150 s).

We next interrogated the viral HA-receptor interaction via two means: by NA inhibition and by sialidase treatment (29). In the control and oseltamivir (20 µM)-treated MDCK cells, the number of bound IAV-AF594 virions per AFM field of view was 4.1 ± 1.5 and 3.6 ± 1.5, respectively (**Fig. S6**). Cells treated with 0.65, 2.5, 10 mUnits/mL sialidase at 37°C for 30 min, bound 3.8 ± 2.1, 1.4 ± 0.9, and 0.1 ± 0.1 virions per AFM field of view, respectively (**Fig. S6**). We selected 2.5 mUnits/mL sialidase treatment for further experiments. The cell surface diffusion of the IAV particles was detected by ViViD-AFM and the trajectory of the IAV virions and their diffusion log_10_D are shown in Fig. 2D, E. The diffusion log_10_D showed a Gaussian distribution (**Fig. 2E**). In the control cells the log_10_D was −2.82 ± 0.27 log_10_ μm^2^/s (N=40), whereas the sialidase-, oseltamivir (20 µM)-, zanamivir (20 µM)-, and zanamivir (30 µM)-treated groups showed values of −2.32 ± 0.39 (N=17), −3.10 ± 0.31 (N=28), −2.88 ± 0.23 (N=30), and −3.28 ± 0.22 (N=28) log_10_ μm^2^/s, respectively (**Fig. 2E**). The increase in the log_10_D values in the sialidase-treated group (**Fig. 2E**) was consistent with in vitro findings of the inverse correlation between diffusion coefficient and sialic acid density (28). Compared with the control, a significant decrease in IAV diffusion was observed in the oseltamivir (20 µM)-treated (p=0.007) and zanamivir (30 µM)-treated (p<0.0001) cells (**Fig. 2E, Fig. S7**). Conversely, a significant increase in IAV diffusion was seen in the sialidase-treated (p<0.0001) cells (**Fig. 2E, Fig. S7**).

## IAV cell surface diffusion reduces with the onset of clathrin coat assembly

Cell surface diffusion facilitated the IAV virion to multivalently engage sialylated receptors. Receptor clustering activates signaling and receptor mediated uptake by a clathrin-dependent or -independent route (9). MDCK cells stably expressing clathrin light chain a (CLCa)-EGFP were inoculated with IAV-AF594 at an MOI of 100 at 37°C, followed by ViViD-AFM imaging at 5 s intervals (**Fig. 3A**). In this example, clathrin coat assembly under the virion began 50 s after the start of observation (**Fig. 3A, B, Movie S4**). The diffusion coefficient D of IAV, calculated from the MSD plot, was 0.0035 µm²/s and 0.0001 µm²/s before and after the onset of clathrin coat assembly, respectively (**Fig. 3C, D**). The reduction in the IAV diffusion coefficient D after clathrin coat assembly was also observed under sialidase and oseltamivir (20 µM) treatments (**Fig. 3E**). In control cells, IAV virions took an average of 50 s from the start of observation to trigger clathrin coat assembly (**Fig. S8**). In comparison, sialidase- and oseltamivir (20 µM)-treated cells required 368 s (p=0.0288) and 79 s (p=0.7483), respectively (**Fig. S8**).

**Fig. 3.**
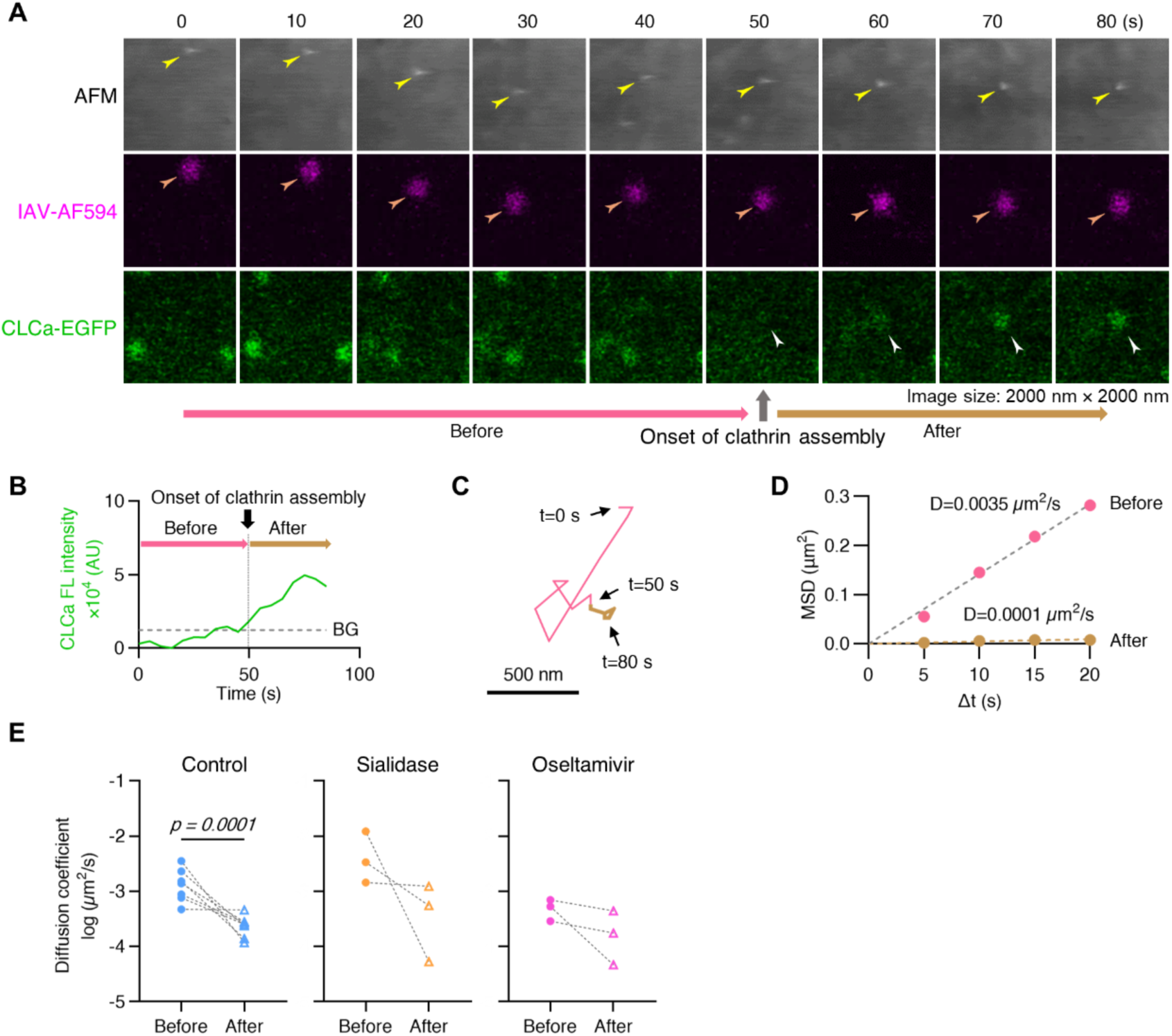
IAV cell surface diffusion before and after clathrin assembly. Live MDCK cells expressing clathrin light chain a (CLCa) fused with EGFP (CLCa-EGFP) were inoculated with IAV-AF594 at 37°C and subjected to 5 s interval time-lapse ViViD-AFM imaging. **(A)** Time-lapse detection of clathrin assembly (CLCa-EGFP, white arrowheads) and diffusing IAV virion (fluorescence, orange arrowheads; morphology, yellow arrowheads) by ViViD-AFM. Image size: 2000×2000 nm^2^. **(B)** Plot of fluorescence (FL) intensity of CLCa at the sites of virion attachment over time. The time when CLCa intensity exceeded the background level (BG) was defined as the onset of clathrin assembly. **(C)** Trajectory of a cell surface IAV virion before and after clathrin assembly, each colored with pink and brown, respectively. Scale bar: 500 nm. **(D)** Quantification of diffusion of a single IAV virion at the cell surface before and after clathrin assembly. Diffusion coefficient (D) is indicated in the graph. MSD, mean squared displacement. **(E)** Dot plot showing diffusion coefficients of the individual IAV virions before and after clathrin assembly under control (N=7), sialidase-treated (N=3), and oseltamivir-treated (N=3) conditions. Plots of the identical virions are connected by dashed lines. AU, arbitrary units.

## Ruffles promote pit closure above the cell membrane baseline during IAV endocytosis

Next, we analyzed IAV internalization via CME, the major uptake route for the virus (9, 11). MDCK cells stably expressing CLCa-EGFP were inoculated with IAV-AF594 at an MOI of 100 at 37°C, followed by ViViD-AFM imaging at 5 s intervals (**Fig. 4A, S9**). The time point at which the virion disappeared from AFM view and was still detectable by fluorescence was defined as t = 0 s (**Fig. 4A, S9**). The onset of clathrin coat assembly preceded membrane ruffle formation (**Fig. 4A, B, C, S9, 10A**), and ruffles approximately 100-nm in height promoted closure of the clathrin-coated pit (CCP) (**Fig. 4A, B, C**). We observed that as the clathrin coat matured, the IAV virion sank into the cell membrane and the height of the virion above the membrane baseline decreased from approximately 85 to 60 nm (**Fig. 4C**). Simultaneously, the membrane ruffle volume increased from ca. 3 × 10^6^ to 8 × 10^6^ nm^3^ and reached a height of over 120-nm above the membrane baseline (**Fig. 4A, B, C, Fig. S10A**). The closure of an IAV-containing CCP typically occurred 50-nm above the baseline of the cell membrane (**Fig. 4C**). Both the clathrin intensity and membrane ruffle volume peaked at t = 0 s (**Fig. 4C, Fig. S10A**). CCP diameter measurements during CME in the absence of IAV showed that clathrin intensity and ruffle volume peaked when the CCP closed (**Fig. S10B, C, D**). The presence of the IAV virion during CME, however, had a major impact on the membrane ruffle size. The average ruffle volume during IAV virion (+) CME and IAV virion (-) CME was 5.9 × 10^6^ compared to 3.2 × 10^6^ nm^3^ (p=0.0137) (**Fig. 4D, Fig. S11A**). Sialidase or oseltamivir treatment did not affect ruffle size during CME of IAV virions (**Fig. S11B**).

**Fig. 4.**
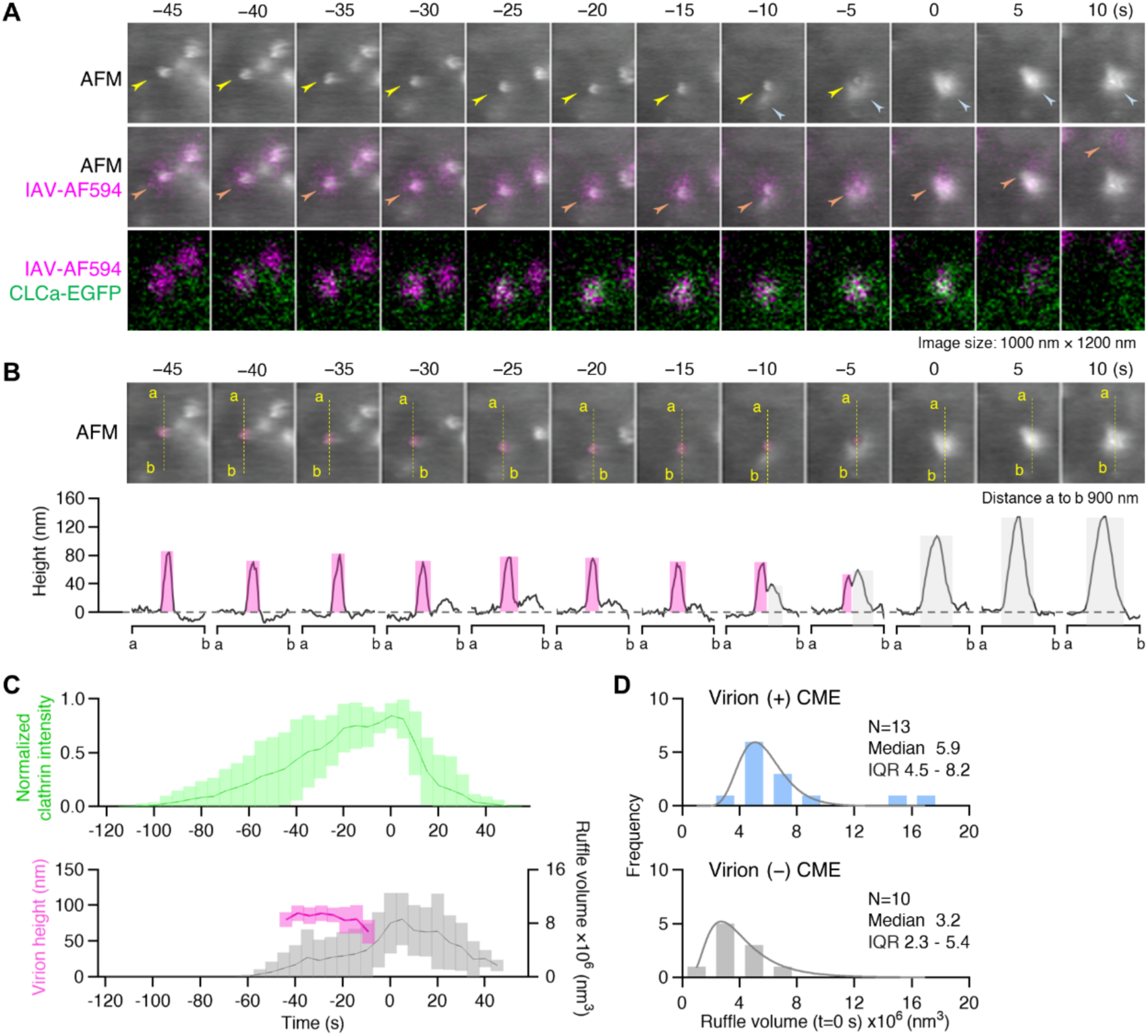
Membrane ruffles promote IAV clathrin-mediated endocytosis. MDCK cells expressing CLCa-EGFP were inoculated with IAV-AF594 and imaged by ViViD-AFM at 37°C at 5 s intervals. **(A)** Time-lapse imaging of IAV internalization. Top to bottom: sequential images of AFM, the merge of AFM and IAV-AF594 (magenta), and the merge of IAV-AF594 and CLCa-EGFP (green). Arrowheads: virion morphology (yellow), membrane ruffles (blue) and virion fluorescence (orange). Virion morphology disappeared at t=0 s. Image size: 1000 × 1200 nm^2^. **(B)** Cross-sectional profiles (bottom) along dashed lines over the virion (magenta) in AFM images (top) of panel (A). Virions and membrane ruffles in profiles are colored with magenta and gray, respectively. Gray dashed line: membrane baseline (height, 0 nm). **(C)** Clathrin intensity (top, green) (N=10), virion height (bottom, pink) (N=7) and ruffle size (bottom, gray) (N=10) during IAV endocytosis. Data are mean ± standard deviation. **(D)** Comparison of the membrane ruffle volume between virion (+) (top) (N=13) and virion (-) CME (bottom) (N=10). Each histogram fitted using a normal distribution in logarithm. IQR, interquartile range.

An interesting feature of membrane ruffles during CME was their size heterogeneity. In the presence of an IAV virion, CCP closure was accompanied by large (>8 × 10^6^ nm^3^) and medium (4 - 8 × 10^6^ nm^3^) sized ruffles that persisted for up to more than 15 s after CCP closure (**Fig. S12**). In the absence of a virion, however, ruffles were either medium or small (<4 × 10^6^ nm^3^) (**Fig. S12**). These observations confirmed that IAV-containing CCPs required significantly larger membrane ruffles to complete CME. In some cases, IAV macropinocytosed by very large membrane ruffles (**Fig. S13, Movie S5**) (2, 11, 30), and a strong correlation between the height of ruffles and the EGFP-Lifeact fluorescence intensity was observed (**Fig. S13**).

In a separate ViViD-AFM experiment in which IAV CME was completed by a membrane ruffle higher than 100 nm (**Fig. S14A, B, C, Movie S6**), we observed that clathrin coat assembly and membrane ruffle formation occurred multiple times prior to internalization (**Fig. S14C, Movie S7**). Interestingly, not only did the membrane ruffles ‘chase’ the virion, but their size also increased as the number of failed uptake events increased (**Fig. S14C, Movie S7**). During its interactions with membrane ruffles, the virion diffused on the cell membrane undergoing several cycles of clathrin coat assembly and disassembly before endocytosis (**Fig. S14C, Movie S7**). Following IAV endocytosis, microtubule-based directed motion of the endosome increased the intracellular virion velocity (µm/s), as reported previously (9, 29) (**Fig. S14D, Movie S8**).

Clinical IAV isolates are filamentous and can reach up to 30 µm in length (31). Filamentous IAV enters cells by macropinocytosis, and the low pH in late endosomes leads to a fragmentation of the filamentous virions via a conformational change in the viral M2 (32). To analyze the interplay between filamentous IAV virions and the cell membrane, we used the influenza virus A/Udorn/307/72 (H3N2) strain labeled with DiI (IAV-DiI) (**Fig. 5**). Glass-bound IAV-DiI imaged by ViViD-AFM showed linear filaments that showed a mean value of 79.3 ± 5.0 nm in height and 1425 ± 447.3 nm in length (N=7) (**Fig. 5A, B**).

**Fig. 5.**
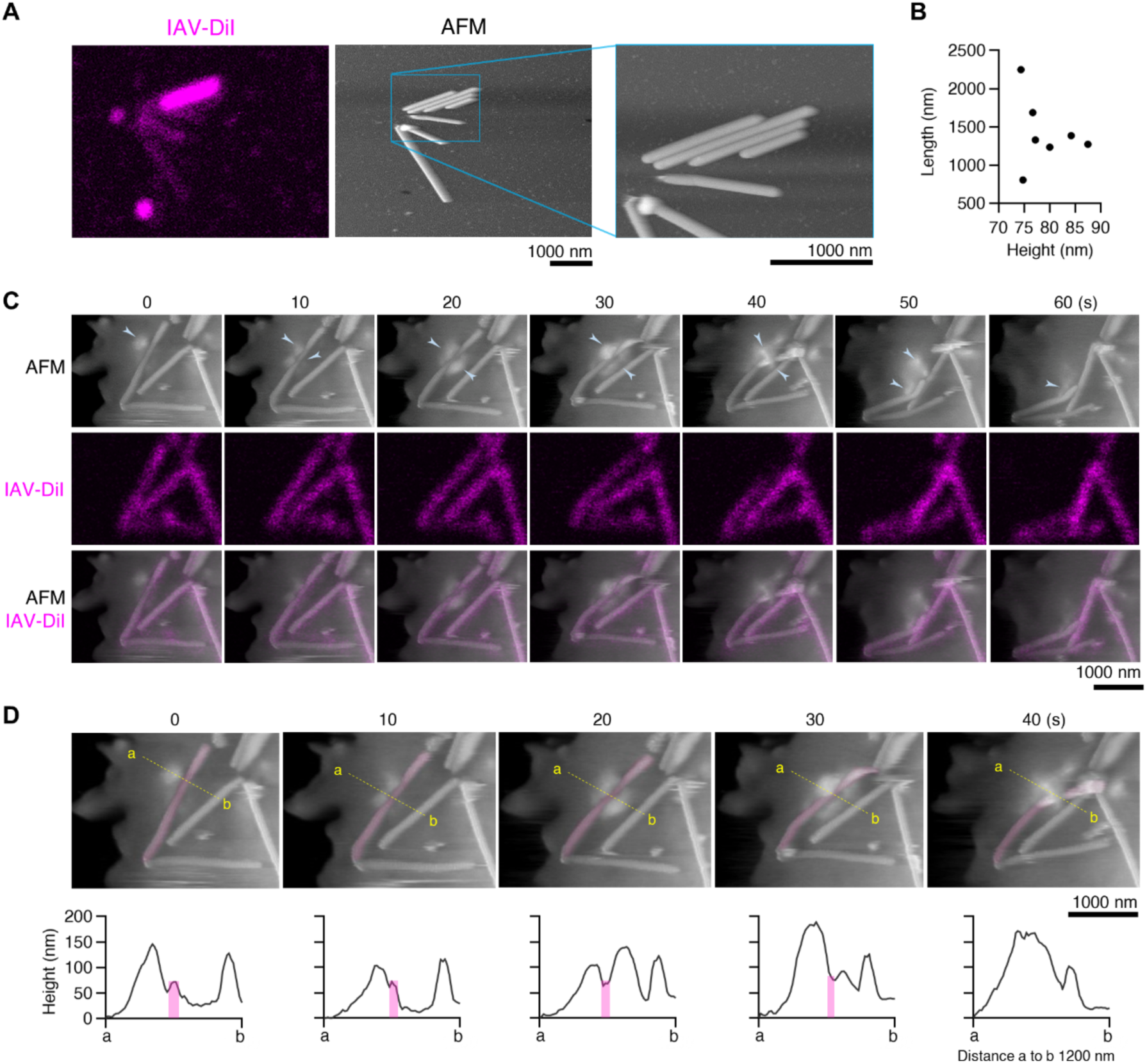
Morphological deformation of filamentous IAV virions by membrane ruffles. **(A)** Fluorescence (left) and morphology image (right) of glass-bound IAV Udorn virions labeled with DiI (IAV-DiI) acquired by ViViD-AFM. **(B)** Dot plot of height (x) and length (y) of IAV-DiI virions in panel (A) (N=7). **(C)** Time-lapse ViViD-AFM imaging of filamentous IAV-membrane ruffle interplay. MDCK cells were inoculated with IAV-DiI and imaged by ViViD-AFM at 27°C at 10 s intervals. Top to bottom: sequential images of AFM, IAV-DiI (magenta) and the merge of AFM and IAV-DiI. Arrowheads: membrane ruffles. **(D)** Cross-sectional profiles (bottom) along dashed lines over the virion in AFM images (top) of panel (C). The virion interacting with membrane ruffles are colored with magenta. (A-D) Scale bars: 1000 nm.

MDCK cells were inoculated with IAV-DiI and imaged by ViViD-AFM at 27°C at 10 s intervals. We found that membrane ruffles 100-200-nm in height appeared on both sides of a filamentous virion (**Fig. 5C, D, Movie S9**). The membrane ruffles induced IAV filament fragmentation and a 428 nm-long fragment including the virion tip detached from the main body (**Fig. 5C, D, Movie S9**). Next, MDCK cells inoculated with IAV-DiI were imaged by confocal microscopy at 27°C at 10 s intervals (**Fig. S15, Movie S10**). An IAV filament on the cell surface deformed over time until the tip of the filament ca. 200-250 nm in length detached from the main body, followed by internalization and intracellular trafficking by directed motion (**Fig. S15, Movie S10**). Taken together, these observations suggest that ruffle-mediated fragmentation of filamentous IAV virions promotes host cell entry (**Fig. 5C, D, S15, Movie S9, S10**).

## Discussion

In this study, we enhanced the minimally invasive capabilities of AFM by using an ultra-narrow cantilever that preserves the interaction between IAV virions and the plasma membrane. Utilizing live-cell correlative imaging of morphology and fluorescence with ViViD-AFM, we thoroughly analyzed the early stages of IAV-cell membrane interplay during the early stages of virus host cell entry. ViViD-AFM elucidated the diffusion characteristics of spherical IAV virions that lead to endocytosis via CME, the viral and cellular factors influencing IAV diffusion, and the actin-regulated cell membrane dynamics that promote endocytosis of spherical and filamentous IAV virions (**Fig.S16**).

Diffusion is essential for an incoming virion to engage enough cell surface receptors for clustering and signaling. Higher sialic acid density correlates with a lower diffusion coefficient (28), suggesting that the heterogeneous density of sialic acids on the cell surface accounts for the varying diffusion coefficients of IAV (**Fig. 2B, C, E**). Previous studies have shown that efficient virion uptake requires a balance between viral HA-receptor binding and neuraminidase activity (26, 27, 33–35). There were two distinct steps in the IAV diffusion process as diffusion dropped significantly after onset of clathrin coat assembly (**Fig. 3**).

ViViD-AFM also revealed that actin-rich membrane ruffles promoted CCP closure during spherical IAV CME. This occurred ca. 50-nm above the cell membrane baseline (**Fig. 4A, B, C, S14A, B**). Akin to observations where virion length dictated the requirement for actin assembly during CME of vesicular stomatitis virus (36), IAV CME required significantly larger ruffles than CME without virus (**Fig. 4C, D, S10, S11, S12**). Clinical influenza viruses are filamentous and can reach up to 30 µm in length. Filamentous virions are thought to enter cells by micropinocytosis followed by low pH-induced fragmentation in endosomes (32). Our observations suggest a mechanism of fragmentation via membrane ruffles at the cell surface (**Fig. 5C, D, S15, Movie S9, S10**). This mechanism may be a viral cue that facilitates cell uptake of filamentous IAV by reducing its size (37).

Our findings using ViViD-AFM provide new insights into the importance of the dynamics of the actin cytoskeleton in the early steps of IAV host cell entry. We propose a model in which following attachment to sialic acid receptors on the cell surface, spherical IAV virions diffuse laterally with the help of viral neuraminidase activity (**Fig. S16A**) (23–27). Being multivalent, viruses usually bind to multiple receptors. Following receptor clustering and reduced diffusion, the IAV virion primes the cell for entry by activating cell signaling pathways, which typically involve receptor tyrosine kinases (2). Clathrin coats are assembled around the virion (9) and dynamics of the actin cytoskeleton generate membrane ruffles in the vicinity. The virion sinks slowly into the cell membrane, and medium to large membrane ruffles engulf the virion before it sinks completely into the CCP (**Fig. S16B**). Following dynamin-mediated fission and clathrin-coated vesicle formation, the internalized virion is directed to early endosomes and transported towards the nucleus along microtubules. After attachment to the cell surface, filamentous IAV virions undergo fragmentation by membrane ruffles from both sides (**Fig. S16C**) which reduces their size and facilitates internalization (30, 32).

In summary, our ViViD-AFM imaging system has the potential to significantly advance our understanding of the mechanobiology of pathogen-membrane interactions. In the future, ViViD-AFM could be applied to study various biological activities occurring at the cell surface, such as the formation and uptake of spherical and filamentous IAVs, extracellular vesicles (EVs) and lipid nanoparticles (LNPs) (38, 39). EVs and LNPs are used in drug delivery to enhance therapeutic efficacy (38, 39), and a critical aspect of EV drug delivery is its cellular uptake, with EVs diffusing on cell surfaces (40) and endocytosing (41). ViViD-AFM therefore is a powerful tool that opens new avenues of research in membrane biology, virus-host interactions, and drug discovery.

## Acknowledgments

We thank Nanoworld AG, NanoTools GmbH, and Mr. Nobuhiro Saito from NanoAndMore Japan for providing ultra-narrow cantilevers. We also extend our gratitude to Dr. Yuki Suzuki for reading the manuscript and Ms. Chiharu Sakai for her technical support. Our thanks go to Eri Komatsu from FEI Company Japan Ltd. for acquiring SEM images. Additionally, we appreciate the technical assistance from Ms. Yuka Imaoka, Mr. Akira Yagi, and Mr. Shuichi Ito of Olympus Corporation in developing instruments. We are grateful to Dr. Kazuhiro Aoki for providing pCX4puro-CRY2-cRaf and to Dr. Asuka Nanbo for supplying eGFP-CLCa. Dr. Yusuke Ohba is acknowledged for providing laboratory facilities for our experiments.

## Author contributions

Conceptualization of the methodology for viral entry: MB and YY. Design of the AFM methodology: YU and NS. Design of the experiments for viral entry: AY, TS, NS, and YY. Investigation: AY, YU, TS, and NS. Visualization: AY, YU, NS, and YY. Funding acquisition: AY and YY. Project administration: NS and YY. Writing – original draft: AY, YU, NS, and YY. Writing – review & editing: AY, YU, TS, MB, NS, and YY.

## Materials and Methods

### Establishment of a minimally invasive AFM cantilever

At room temperature, the force of interaction between the virion and the plasma membrane of MDCK cells is approximately 10-25 pN (22). To reduce the potential mechanical effect on virus-cell interactions, a new cantilever with reduced spring constant was designed. The spring constant (k_c_) and resonance frequency (f_c_) of a cantilever are related to its shape (length l, width w, thickness t) as follows:

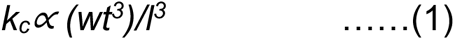

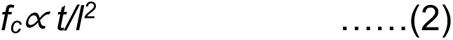

Based on these equations, we employed the ultra-narrow cantilever which has a length (l) of 9.0 μm, width (w) of 0.8 μm, and thickness (t) of 0.1 μm (Fig. 1B). The new ultra-narrow cantilever has a spring constant of 0.04 N/m, and a resonance frequency in liquid over 300 kHz. This design reduced the minimum peak force to 10 pN, which is small enough to preserve virus-cell surface interactions.

### Preparation of fluorescently labeled viruses

The influenza A/WSN was prepared as previously described (42). Briefly, the virus was propagated in 30 chicken eggs at 35°C for 2 days. After one day of incubation at 4°C, the allantoic fluid was collected and clarified by centrifugation at 10,000 rpm for 20 min. The clarified fluid was then concentrated by centrifugation at 100,000 × g for 90 min. Further purification involved two cycles of centrifugation using Iodixanol (Sigma-Aldrich, St. Louis, MO, USA) gradient ranging from 10 to 40% at 100,000 × g for 90 min. Viral bands were collected and resuspended in NTC buffer (100 mM NaCl and 20 mM Tri-HCl, pH 7.4, 5 mM CaCl^2^). The viral titer was determined to be 1.0 × 10^10^ plaque-forming units per mL in MDCK cells. Aliquots of the virus were stored in NTC buffer at - 80°C until use.

The influenza A/WSN/33 (H1N1) virus was labeled with Alexa Fluor 594-NHS ester dye (A37572, Thermo Fisher, Carlsbad, CA, USA) as follows. NaHCO^3^ (pH ≈ 8) was added to the virus solution to a final concentration of 0.14 M. Alexa Fluor 594 ester dye, suspended in DMSO, was then added to obtain the final dye concentration of 50 µM. The virus, at a concentration of 0.8×10^9^ pfu/ mL, was incubated with dye for 1 hour in the dark at room temperature. To remove unbound dye, the reaction mixture was loaded onto a PD MiniTrapTM G-25 column (28918007, GE Healthcare, Little Chalfont, UK) pre-equilibrated with HBS buffer (150 mM NaCl, 20 mM HEPES). The influenza A virus conjugated with Alexa Fluor 594 (IAV-AF594) was then eluted from the column with HBS buffer.

The influenza A/Udorn307/72 (H3N2) virus labeled with lipophilic carbocyanine dye (DiI, D282, Thermo Fisher) was prepared as follows. MDCK cells were inoculated with the virus at 37°C and incubated for 1 hour. Inoculated cells were then labeled with DiI by incubation with 5 µg/mL dye for 20 minutes. After 2 days of incubation at 37°C, the supernatant was collected and clarified by centrifugation at 2,000 × g for 10 minutes. The viral titer was determined to be 1.0 × 10⁸ PFU/mL in MDCK cells. The virus sample was stored at 4°C until use.

### Plasmids

cDNA for CLCa was amplified from eGFP-CLCa (a kind gift from Dr. Asuka Nanbo, Nagasaki University) (43) by PCR with the primers, CLCa forward (GGCTCGAGATGGCTGAGCTGGATCC) and CLCa reverse (TAGCGGCCGCTCAGTGCACCAGCGGG). The amplified sequence was subcloned into the XhoI/NotI sites of the pCX4puro-EGFP vector (a gift from Dr. Kazuhiro Aoki, National Institutes for Basic Biology). For Lifeact expression, pFX-Lifeact-EGFP was used (44).

### Cell culture

MDCK cells (CRL-34, American Type Culture Collection, Manassas, VA, USA) were cultured under a 5% CO_2_ humidified atmosphere at 37°C in Dulbecco’s Modified Eagle’s Medium (DMEM, Sigma-Aldrich) supplemented with 10% fetal bovine serum (Thermo Fisher Scientific). For transient expression of Lifeact-EGFP, the pFX-Lifeact-EGFP expression vector was transfected into MDCK cells using Polyethylenimine Max (Polysciences, Warrington, PA, USA). To establish cell lines stably expressing CLCa-EGFP, expression vectors for CLCa fused with EGFP were linearized by ScaI and introduced into the MDCK cells using nucleofection according to the manufacturer’s protocols (Amaxa Biosystems, Cologne, Germany). One day after transfection, cells were cultured in DMEM containing 4 µg/mL Puromycin (Wako, Kyoto, Japan). Resistant colonies were collectively isolated two days after transfection. Stably expressing cells were maintained in DMEM supplemented with 10% FBS.

### ViViD-AFM system

The ViViD-AFM composed of a tip-scan type AFM system (BIXAM™; Olympus Corporation, Tokyo, Japan) with an ultra-narrow cantilever and a multi-color confocal laser scanning microscope system (FV1200, Olympus Corporation). AFM Scanning System Software Version 2.0.2.0 (Olympus Corporation) was used for acquiring time-lapse AFM images and for morphological analysis. The AFM used a tapping mode with phase feed-back control, scanning at a frequency of 30 Hz or 60 Hz in the X direction to image one frame in 10 or 5 s, respectively. 320 data points were sampled in the X direction and 240 in the Y direction. AFM images of cells were displayed in a maximum 6 µm × 4.5 µm × 400 nm (X × Y × Z) area with a resolution of 320 × 240 (X × Y) and 256 shades of an 8-bit gray scale (Z) where bright areas are higher and dark areas are lower. An 880 nm Super Luminescent Diode (Hamamatsu Photonics, Japan) was used for the cantilever optical beam deflection (OBD) sensor. All AFM imaging was performed using a cantilever amplitude of < 95% of its free amplitude, with a minimum peak force 10 pN. A customized ultra-narrow cantilever (USC-F0.8-k0.05; Nanoworld Corporation, Neuchâtel, Switzerland) with an electron beam deposited (EBD) tip < 10 nm radius of curvature at the free end was used. The multi-color confocal laser scanning microscope (FV1200, Olympus Corporation) consisted of an electric inverted fluorescence microscopy IX83, two lasers (473 nm and 559 nm), and two fluorescence channels with High sensitivity GaAsp Detectors. Time-lapse fluorescence images in 12.3 µm × 12.3 µm (X × Y) area with a resolution of 512 × 512 (X × Y) were obtained using a 100 × objective under oil emersion, with a 2 μs/pixel scan speed and 10 or 5 s interval. For correlative imaging, the targeted cell was selected using a phase contrast microscope with a 20 × or 60 × objective lens. Using fluorescence observation with confocal microscope and 100 × objective lens, the target area of the cell was placed on the center of the fluorescence image. The AFM probe with autofluorescence was then approached onto the target area, and ViViD-AFM imaging was initiated. Sequential images captured by AFM and confocal microscope were overlaid by using AviUTL (http://spring-fragrance.mints.ne.jp/aviutl/), based on the AFM tip position as previously described (20). A montage of AFM and fluorescence images was trimmed from the stack images of 6.0 × 4.5 or the 12.3 × 12.3 μm^2^ image using the ImageJ software Version 2.14.0/1.54f (National Institutes of Health; https://imagej.nih.gov/ij/).

### Live ViViD-AFM imaging of viruses at the cell surface

For correlative imaging, glass slides (KB218-0A; Toa optical technologies, LTD, Tokyo, Japan), featuring a circular 15 mm diameter cover glass on a water-repellent printed glass slide were used. MDCK cells were seeded on poly-L-lysine-coated (P4707; Sigma-Aldrich) glass slides one day before imaging. Prior to microscopic observation, the medium was replaced with phenol red-free DMEM/F12 (11039-021; Thermo Fisher Scientific) and the cells were inoculated with IAV-AF594 at a multiplicity of infection (MOI) of 100. ViViD-AFM imaging was initiated 5 min after inoculation at room temperature (27°C) or 37°C. The cell periphery was selected as the imaging area. For neuraminidase inhibition, cells and viruses were treated with 20 µM oseltamivir carboxylate (GS-4021; Cayman, Ann Arbor, MI, USA) or 20 µM or 30 µM zanamivir (S3007; Selleck Chemicals LLC, Houston, TX, USA) 30 minutes prior to virus inoculation. To reduce sialic acid density on the cell membrane, cells were treated with 0.625, 2.5 or 10 mUnit/mL sialidase (N7885; Sigma) for 30 minutes at 37°C and were washed twice with DMEM/F12 medium before virus inoculation.

### ViViD-AFM imaging of viruses immobilized on a glass substrate

To analyze virus morphology and fluorescence labelling efficiency, IAV-AF594 were diluted approximately 200-fold with PBS. The virus was then immobilized onto a glass surface precoated with 1.0 mg/mL fetuin (F3004; Sigma-Aldrich) as described previously (25). The immobilized viruses were observed by ViViD-AFM at room temperature.

### Image analysis

To track and analyze IAV cell surface diffusion, virus trajectories were generated with the use of the Multidimensional Image Analysis module of the MetaMorph software (Molecular Devices, San Jose, CA, USA). From fluorescence stack files, fluorescence spots of IAV-AF594 were segmented with adaptive thresholding and by pairing spots in each frame according to proximity and similarity in intensity. The X-Y-coordinates of IAV over time were exported as an excel file. Mean square displacement (MSD) of virus was plotted over time and the diffusion coefficient (D) was calculated by linearly fitting the MSD curve based on the equation below (45), representing two-dimensional free diffusion.

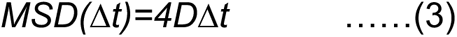

To obtain the fluorescence intensity profile of EGFP-Lifeact or IAV-AF594, the plot profile tool of ImageJ software was used. The cross-section profile of IAV virion and membrane was obtained using AFM Scanning System Software Version 1.6.0.12 (Olympus). From cross-section profiles, pit diameter and virion height were obtained by measuring the distance between 2 points at the edge of the invagination on the section profile as previously described (20) and measuring the distance between the membrane base line and the peak height of virion.

CLCa fluorescence intensity was quantified on with ImageJ by measuring total fluorescence intensity within a 300 nm diameter circle centered at the virion’s center of gravity and subtracting the background from a 1500 nm diameter circle. A moving average plot (2 points) of CLCa fluorescence intensity over time was generated, and the time when CLCa intensity exceeded the background level was defined as the onset of clathrin assembly.

Membrane ruffle volume was measured with ImageJ. On background-subtracted images (using the rolling ball tool), the volume of membrane ruffles was measured within a 500 nm diameter circle centered at the virion’s center of gravity.

### Statistical analysis

Data are presented as mean ± standard deviation (unless indicated otherwise) and were compared between two conditions with Student’s t test or among more than two conditions by Dunnett’s test. The Wilcoxon signed-rank test was applied for comparison between paired samples. P-values are indicated in the respective figure panels. All statistical analyses were performed using Prism 10 Version 10.2.3 (GraphPad Software, Boston, MA, USA).

---

**Fig. S1.**
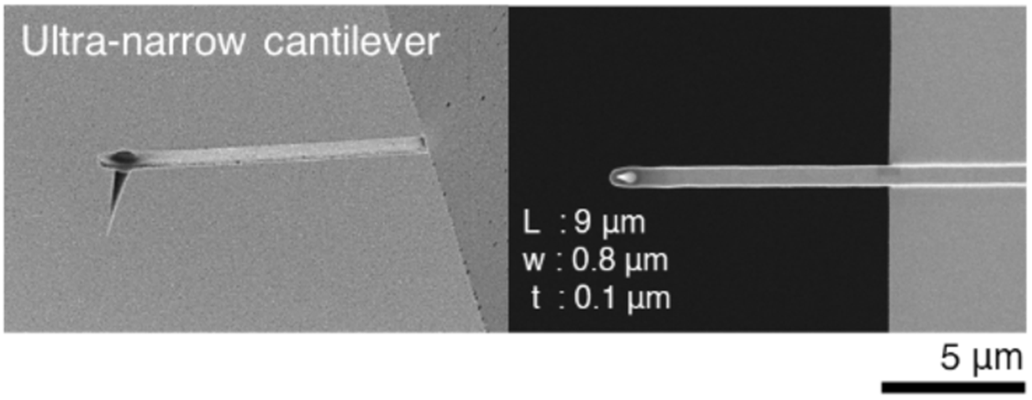
SEM images of the ultra-narrow cantilever for virus-view AFM. Side view (left) and bottom view (right) from the tip side. Scale bar: 5 µm.

**Fig. S2.**
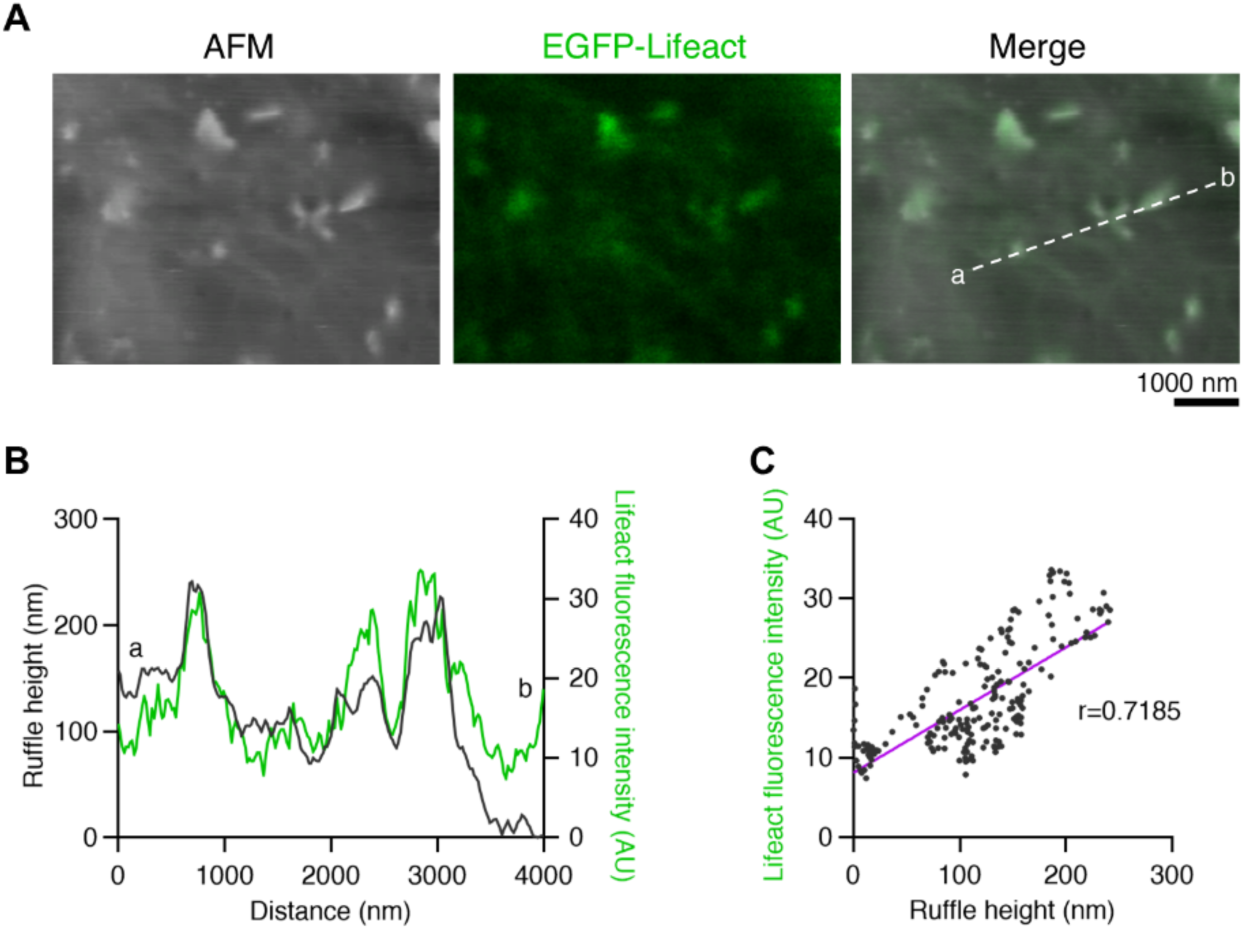
Characterization of MDCK cell surface by ViViD-AFM (related to Fig. 1). Live MDCK cells expressing EGFP-Lifeact were imaged by ViViD-AFM at 27°C at 10 s intervals. **(A)** Representative morphology image of the cell membrane (left), fluorescence image of Lifeact (middle), and merged image (right). Scale bar: 1000 nm. **(B)** Cross-sectional profile of the membrane morphology (gray) and Lifeact fluorescence intensity (green) along the dashed line in panel (A). The lowest height was set as 0 nm. **(C)** Pearson’s correlation coefficient (r) between height of membrane morphology (x axis) and Lifeact fluorescence intensity (y axis), for the profile data in panel (B). AU, arbitrary units.

**Fig. S3.**
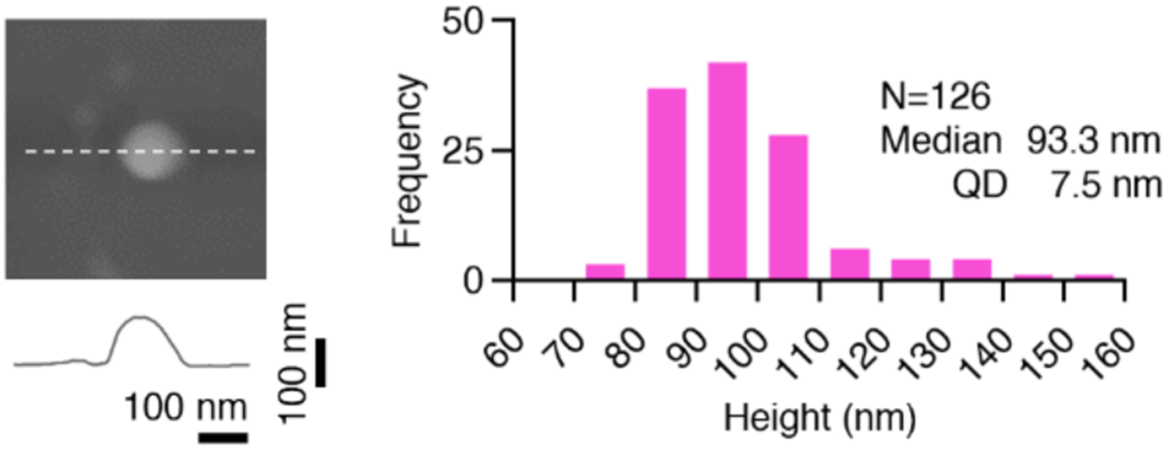
Characterization of fluorescently labeled IAV virions on the glass surface by ViViD-AFM (related to Fig. 1). IAV-AF594 virions were immobilized on fetuin-coated coverslips and subjected to ViViD-AFM at 27°C. Magnified morphology image of a single IAV virion (top left), cross-sectional profile along the dashed line of the virion (bottom left) and histogram of 126 virions heights (right). Scale bars: 100 nm.

**Fig. S4.**
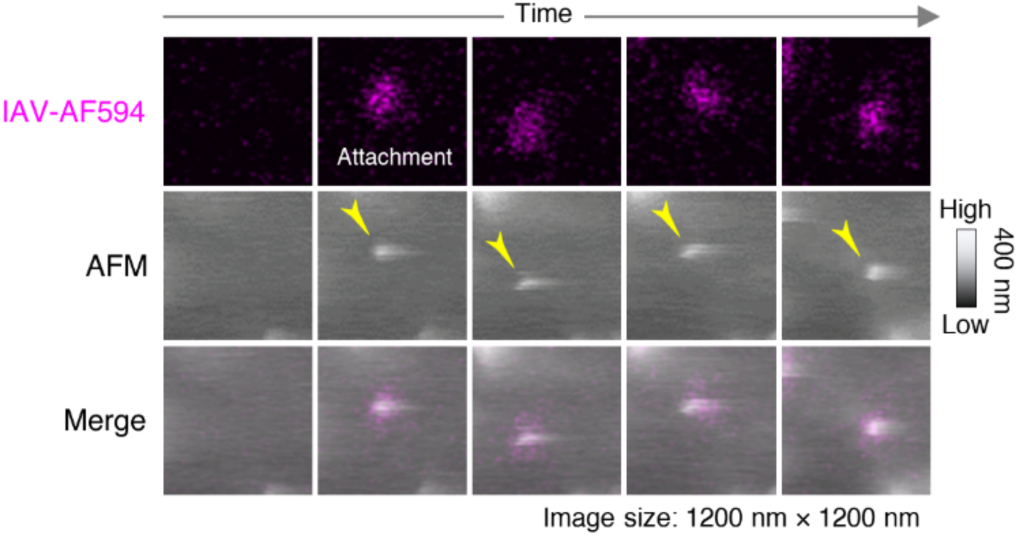
IAV attachment and lateral diffusion observed by ViViD-AFM (related to Fig. 1). Time-lapse imaging of cell surface IAV virions (arrowheads) labeled with Alexa Fluor 594 (IAV-AF594) using ViViD-AFM at 37°C at 5 s intervals. From top to bottom: Fluorescence, morphology, and merged images. The AFM image is a topographic image, with the height direction: bright areas are higher and dark areas are lower. Image size: 1200 ×1200 nm^2^.

**Fig. S5.**
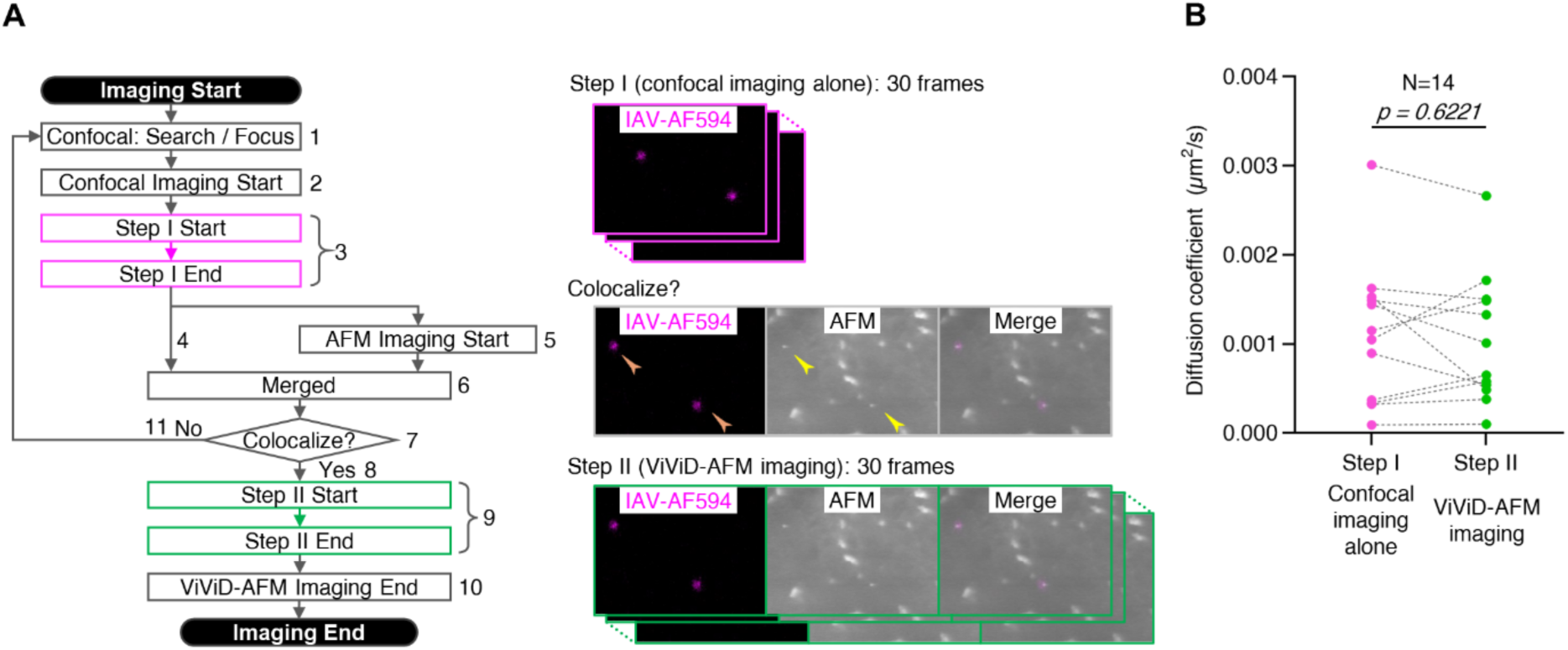
Assessment of the effect of AFM imaging on IAV cell surface diffusion (related to Fig. 1). Identical IAV-AF594 virions were traced for 300 s each for Step I (confocal imaging alone) and Step II (ViViD-AFM imaging). The diffusion coefficient was then determined for steps I and II and was compared to evaluate the effect of AFM imaging. This evaluation was performed on 14 virions. Virions were tracked by time-lapse imaging at 27°C at 10 s intervals. **(A)** Workflow chart to track identical IAV virions in Step I (confocal imaging alone) and the following Step II (ViViD-AFM imaging). Assessment Procedure: (1) First, the confocal microscope was focused on the fluorescent spots (magenta) of the IAV-AF594, which were expected to be at the cell surface. (2) Time-lapse confocal imaging at 27°C at 10 s intervals was started. (3) Thirty frames (300 s) of confocal images were acquired (step I). (4) Confocal imaging was continued after step I. (5) AFM imaging at 10 s intervals was started in parallel with confocal imaging. (6) Confocal and AFM images were overlaid. (7) Virions (yellow arrowheads) that co-localized with fluorescent spots (orange arrowheads) in the merged image were checked. (8) If the virion co-localization with fluorescent spots was detected in (7), ViViD-AFM imaging was continued. (9) Thirty frames (300 s) of AFM and confocal images were acquired (step II). (10) ViViD-AFM imaging was terminated. (11) If virions co-localized with fluorescent spots were not identified in (7), the fluorescent spots may be intracellular IAVs, or the focus of the confocal microscope may not be properly on the cell surface, so either the observation area was changed, or the focus of the confocal microscope was readjusted and step I was started again. **(B)** Dot plot of diffusion coefficients of the IAV virions at the cell surface (N=14) for steps I and II. Plots of the identical virions were connected by dashed lines.

**Fig. S6.**
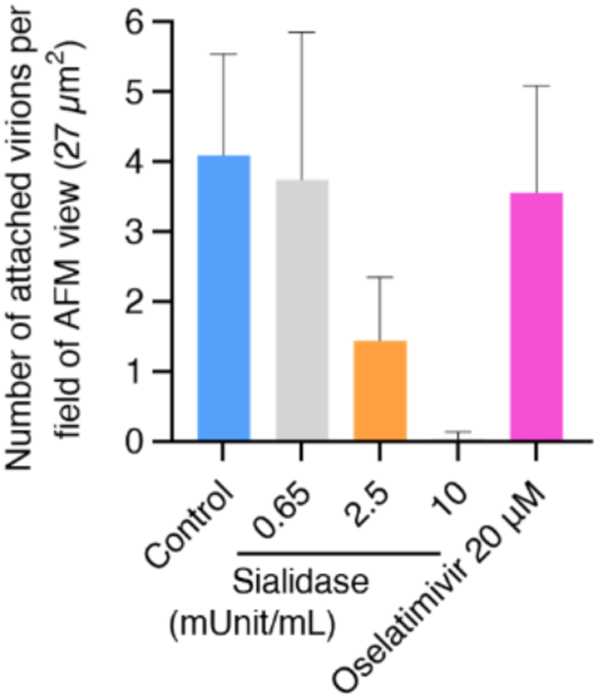
IAV attachment in inhibitor-treated cells (related to Fig. 2). Number of IAV-AF594 virions adsorbed on the cell surface under Control, Sialidase (0.65, 2.5, 10 mUnit/mL)-, and 20 µM Oseltamivir-treated conditions, counted within the AFM observation area (27 µm²) 5 min after inoculation at 27°C. Data are mean ± standard deviation.

**Fig. S7.**
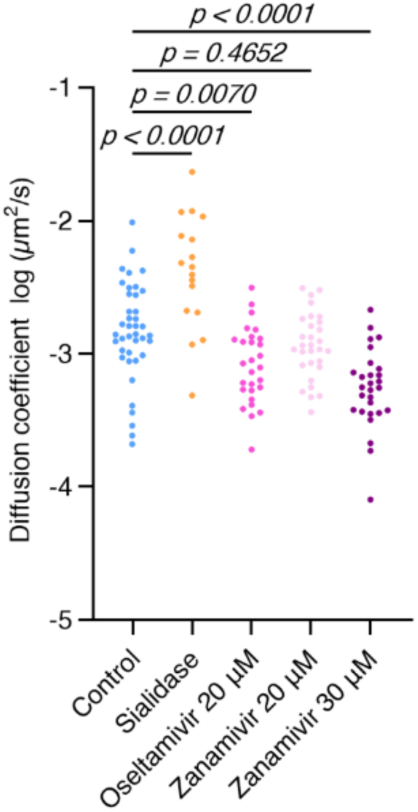
Effect of inhibitors on IAV cell surface diffusion (related to Fig. 2B). Diffusion coefficients of cell surface IAV virions under Control (N=40), Sialidase (=17), 20 µM Oseltamivir (N=28), and 20 µM Zanamivir (N=30) and 30 µM Zanamivir (N=28) conditions, using the same data from Fig. 2B.

**Fig. S8.**
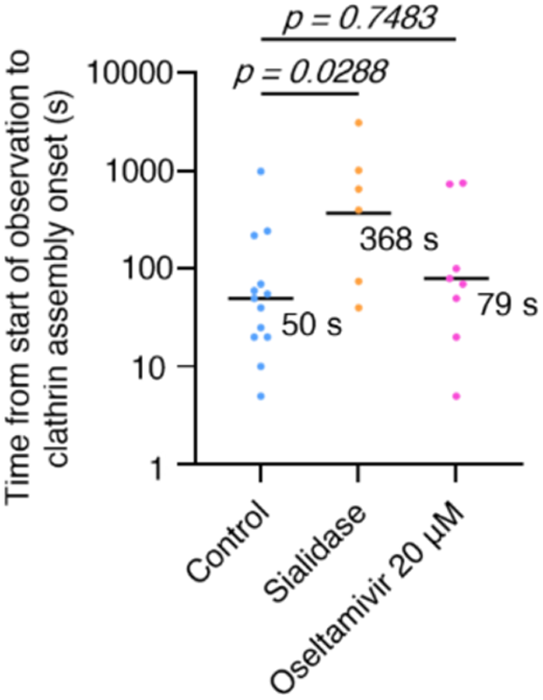
Duration from the start of observation to clathrin assembly onset (related to Fig. 3). Mean values in Control (N=13), Sialidase-treated (N=5), and Oseltamivir-treated (N=8) conditions are shown.

**Fig. S9.**
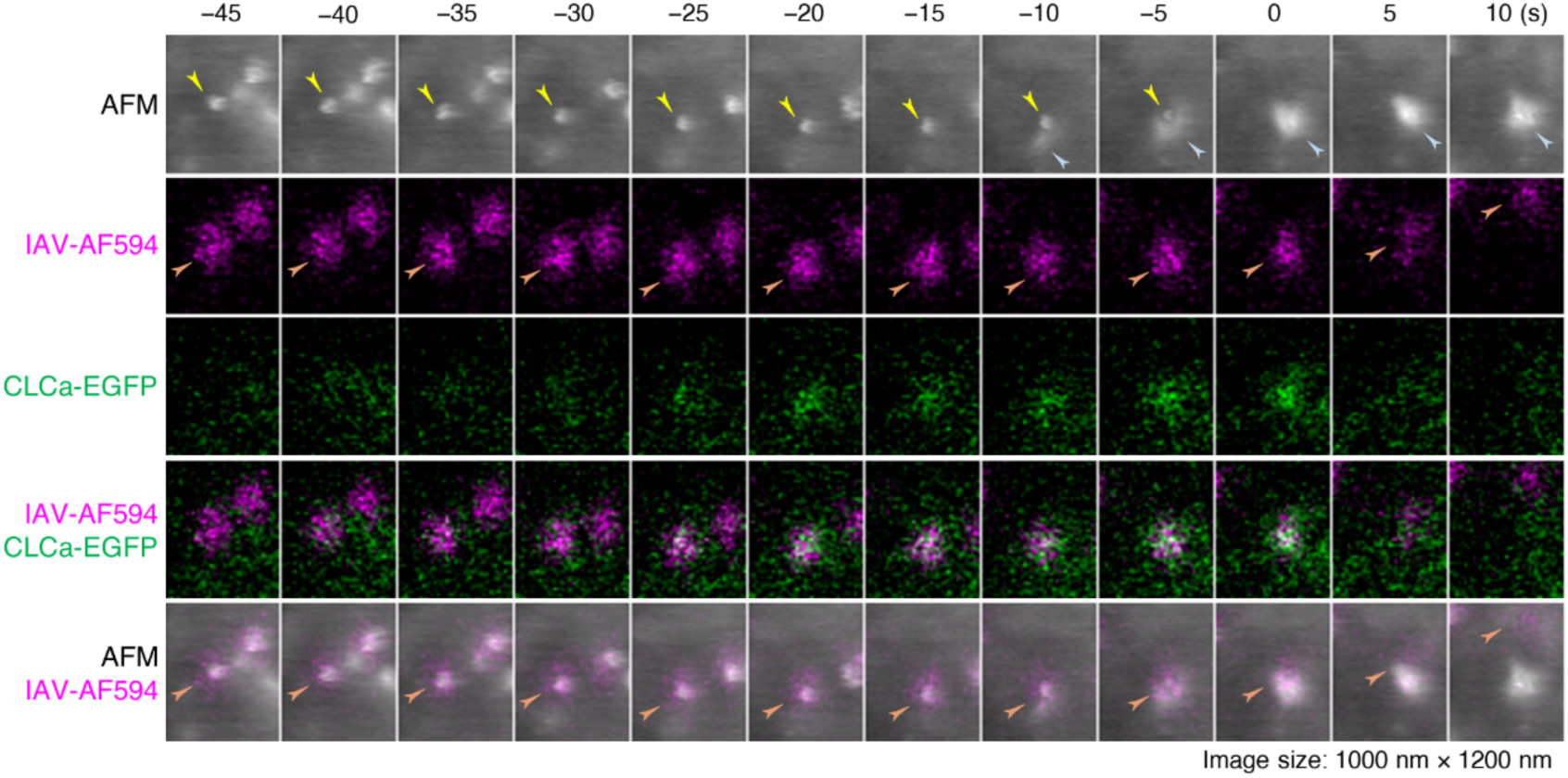
IAV clathrin-mediated endocytosis observed by ViViD-AFM (related to 4A). Live MDCK cells stably expressing CLCa-EGFP were inoculated with IAV-AF594 at 37°C and subjected to 5 s interval time-lapse ViViD-AFM imaging. From top to bottom: sequential images of AFM, IAV-AF594 (magenta), CLCa-EGFP (green), the merge of IAV-AF594 and CLCa-EGFP, and the merge of AFM and IAV-AF594. Arrowheads: virion morphology (yellow), membrane ruffles (blue) and virion fluorescence (orange). t=0 s was set as the moment when the virion morphology was no longer detected. Image size: 1000 × 1200 nm^2^.

**Fig. S10.**
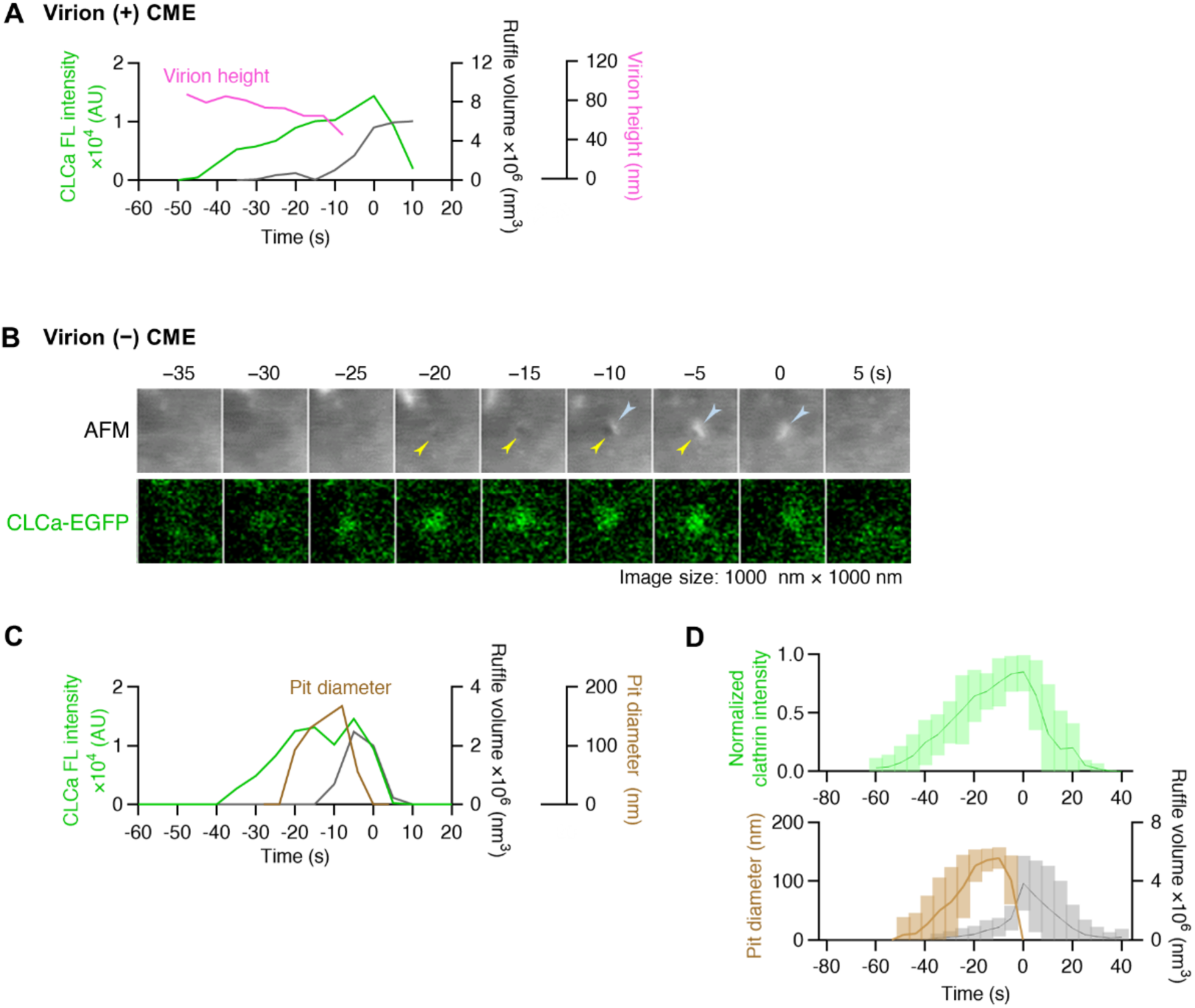
Characterization of clathrin assembly and membrane ruffles during CME with and without IAV using ViViD-AFM (related to Fig. 4). **(A)** Clathrin assembly, membrane ruffle formation, and virion height changes over time in Fig. 4A and B. **(B)** Time-lapse imaging of clathrin-coated pit dynamics during virion (-) CME, acquired in the same cell as Fig. 4A and B. Sequential images of AFM (top) and CLCa-EGFP (green, bottom) are displayed every 5 s. Arrowheads: pit morphology (yellow) and ruffle morphology (blue). t = 0 s was set as the moment when the pit morphology was no longer detected. Image size: 1000 × 1000 nm^2^. **(C)** Clathrin assembly, membrane ruffle, and pit formation over time in panel (B). **(D)** From time-lapse ViViD-AFM data sets of 10 virion (-) CME events, including the one in panel (B), the fluorescence intensity of CLCa-EGFP (top), pit diameter and membrane ruffle volume (bottom) were analyzed and plotted over time. Data are mean ± standard deviation.

**Fig. S11.**
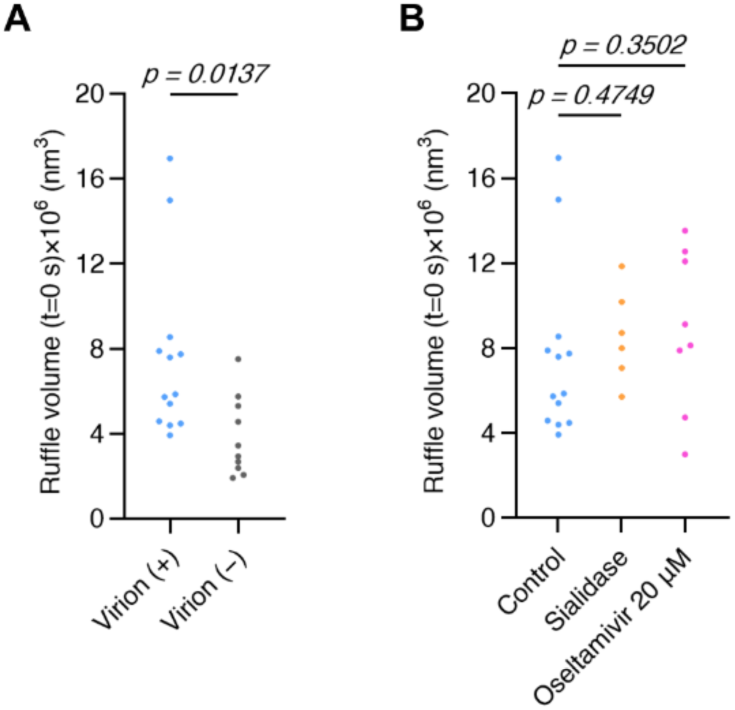
Membrane ruffle size during IAV CME under different conditions (related to Fig. 4). **(A)** Volume of membrane ruffles at t = 0 s in IAV virion (+) (N=13) and IAV virion (-) CME (N=10), using the same data from Fig. 4D. **(B)** Volume of membrane ruffles engulfing IAV virions at t = 0 s under Control (N=13), Sialidase (N=6) and Oseltamivir-treated (N=8) conditions.

**Fig. S12.**
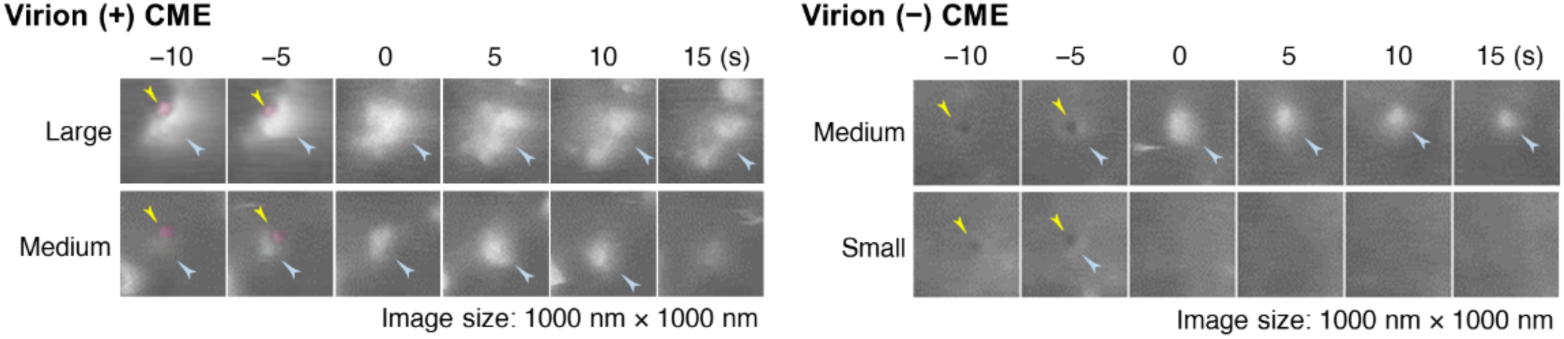
Membrane ruffles with different sizes in IAV virion (+) (left) and virion (-) (right) CME (related to Fig. 4). MDCK cells expressing CLCa-EGFP were inoculated with IAV-AF594 and imaged by ViViD-AFM at 37°C at 5 s intervals. t = 0 s was set as the moment when the virion morphology (magenta) or pit morphology, indicated with yellow arrowheads, was no longer detected. Sequential AFM images show membrane ruffles (blue arrowheads), categorized as large, medium or small according to the ruffle volume at t=0 s. Image size: 1000 × 1000 nm^2^.

**Fig. S13.**
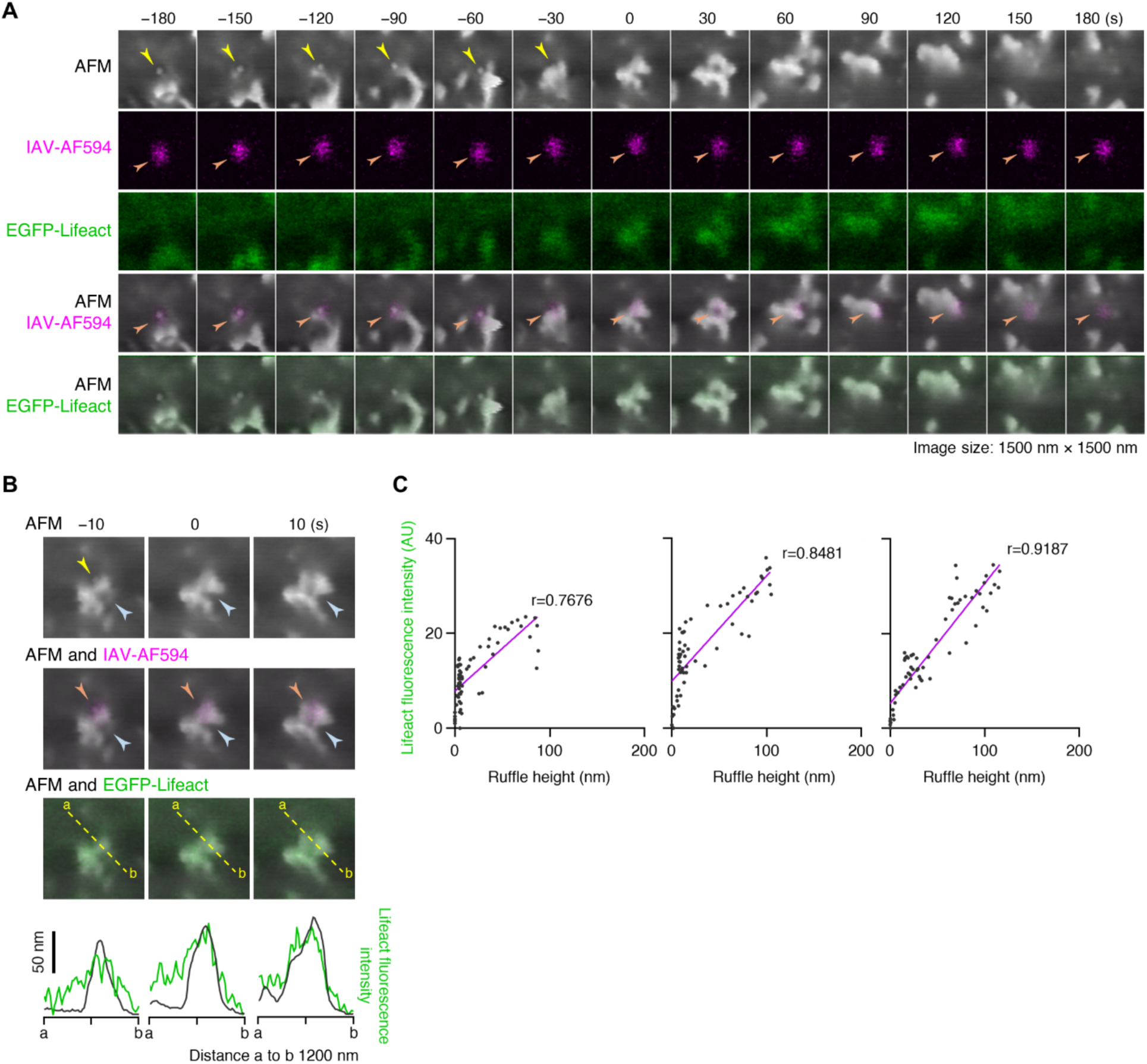
IAV internalization by actin-rich membrane ruffles (related to Fig. 4). MDCK cells transiently expressing EGFP-Lifeact were inoculated with IAV-AF594 at 27°C and subjected to 10 s interval time-lapse ViViD-AFM imaging. **(A)** Live imaging of IAV internalization by ViViD-AFM. From top to bottom, sequential images of AFM, IAV-AF594 (magenta), EGFP-Lifeact (green), the merge of AFM and IAV-AF594, and the merge of AFM and EGFP-Lifeact are displayed every 30 s. t = 0 s was set as the moment when the virion morphology was no longer detected. Image size: 1500 × 1500 nm^2^. **(B)** Correlation analysis of membrane morphology and fluorescence, related to panel (A). From top, sequential images of AFM, the merge of AFM and IAV-AF594 (magenta), and the merge of AFM and the EGFP-Lifeact (green) at t = −10 s, 0 s, and 10 s are displayed. The lowest panel is a plot of the cross-sectional profile of membrane morphology and Lifeact fluorescence intensity along the dashed line in the above merged image. (A, B) Arrowheads: virion morphology (yellow), membrane ruffles (blue) and virion fluorescence (orange). **(C)** Pearson’s correlation coefficient (r) between height of membrane morphology (x axis) and Lifeact fluorescence intensity (y axis), for the profile data in panel (B).

**Fig. S14.**
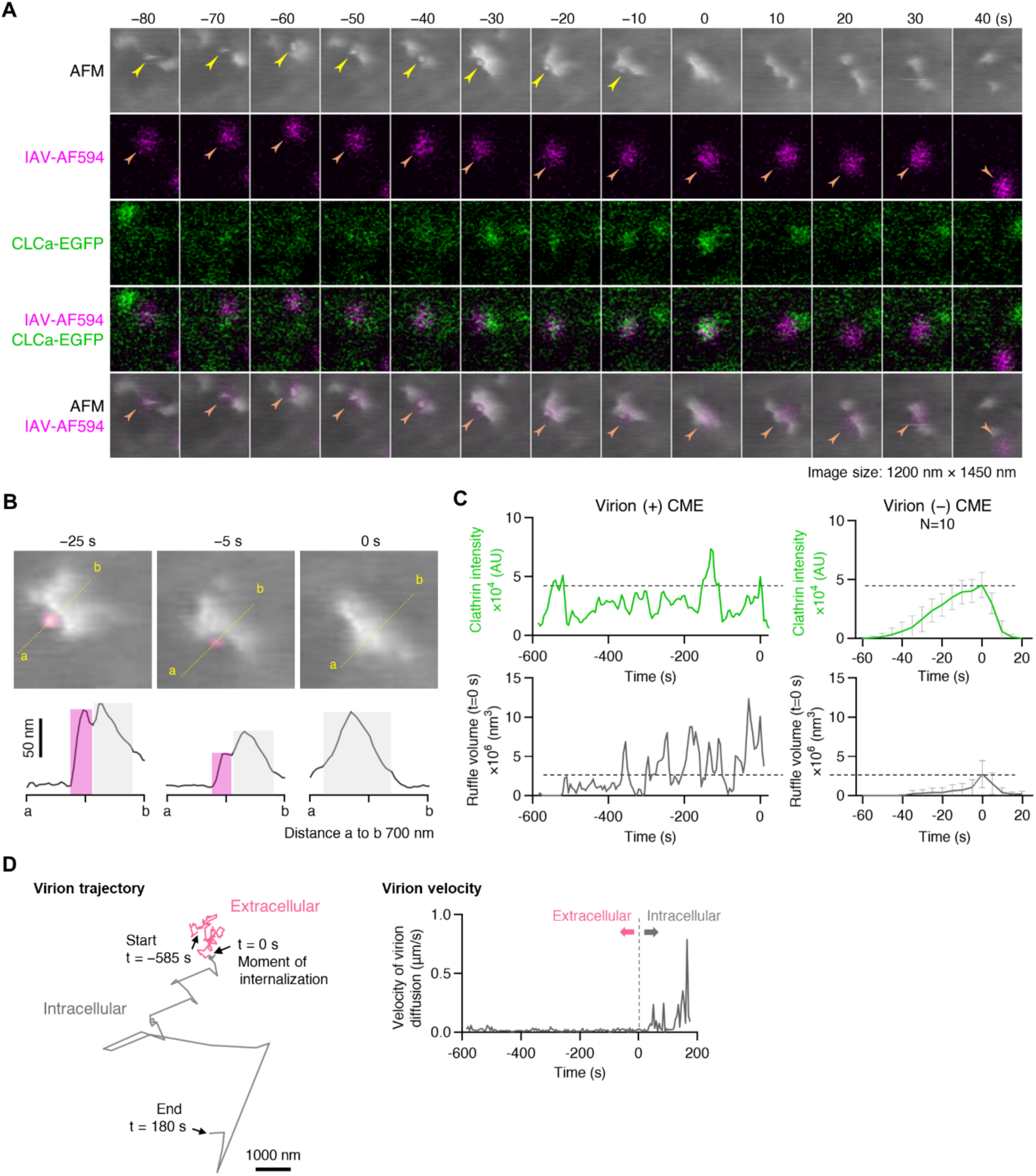
CME of IAV preceded by multiple cycles of clathrin assembly (related to Fig. 4). **(A)** Live imaging of IAV internalization by ViViD-AFM at 37°C at 5 s intervals. From top to bottom, sequential images of AFM, IAV-AF594 (magenta), CLCa-EGFP (green), the merge of IAV-AF594 and CLCa-EGFP, and the merge of AFM and IAV-AF594 are displayed every 10 s. Virion morphology visible in the AFM images is indicated by arrowheads. t = 0 s was set as the moment when the virion morphology was no longer detected. Arrowheads: virion morphology (yellow) and fluorescence (orange). Image size: 1200 × 1450 nm^2^. **(B)** Cross-sectional profiles (bottom) along dashed lines over virion (magenta) in AFM images (top) of panel (A). Virions and membrane ruffles in profiles are colored with magenta and gray, respectively. Scale bar: 50 nm. **(C)** Comparison of clathrin assembly (top, green) and membrane ruffle size (bottom, gray) in virion (+) (left) and virion (-) CME (right) in the identical cell. From IAV endocytosis data in panel (A) and 10 virion (-) endocytosis data sets, mean and standard deviation were calculated and plotted in left and right, respectively. Dashed lines: mean value of maximum clathrin intensity and ruffle size in virion (-) CME (N=10). **(D)** Trajectory of a single IAV virion in extracellular (from t = −585 s to t = 0 s, pink) and intracellular environment (from t = 0 s to t = 180 s, gray).

**Fig. S15.**
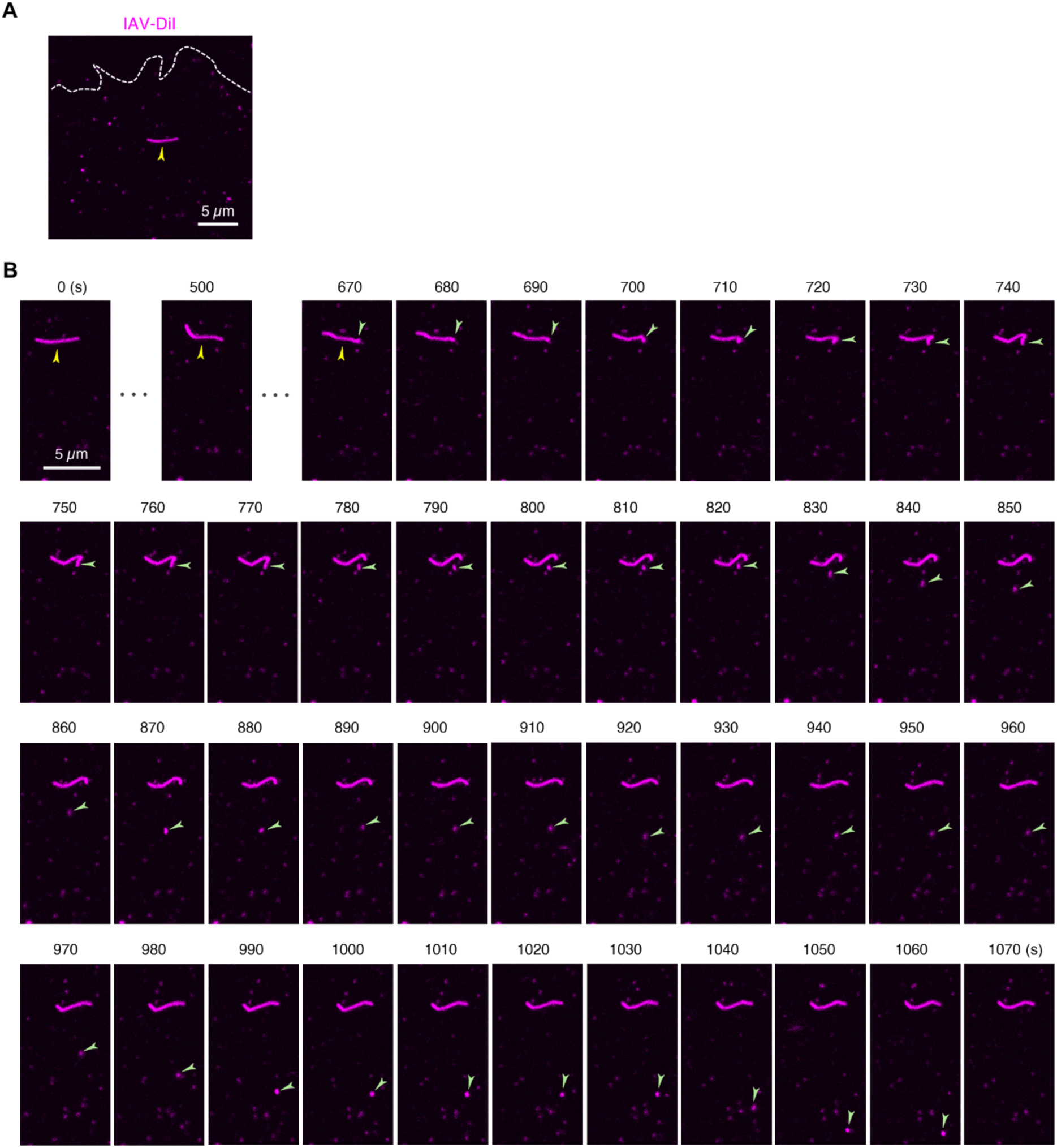
Fragmentation of filamentous IAV followed by transport (related to Fig. 5). MDCK cells were inoculated with IAV-DiI and imaged by confocal microscope at 27°C. **(A)** Filamentous IAV-DiI (arrowhead) imaged by confocal imaging in MDCK cells. Dashed line: cell contour. **(B)** Morphological deformation of filamentous virion in panel (A) captured by time-lapse confocal imaging at 27°C at 10 s intervals. Arrowheads: filamentous virion (yellow), the edge of the filament moving away from the main filament (green). (A, B) Scale bars: 5 µm.

**Fig. S16.**
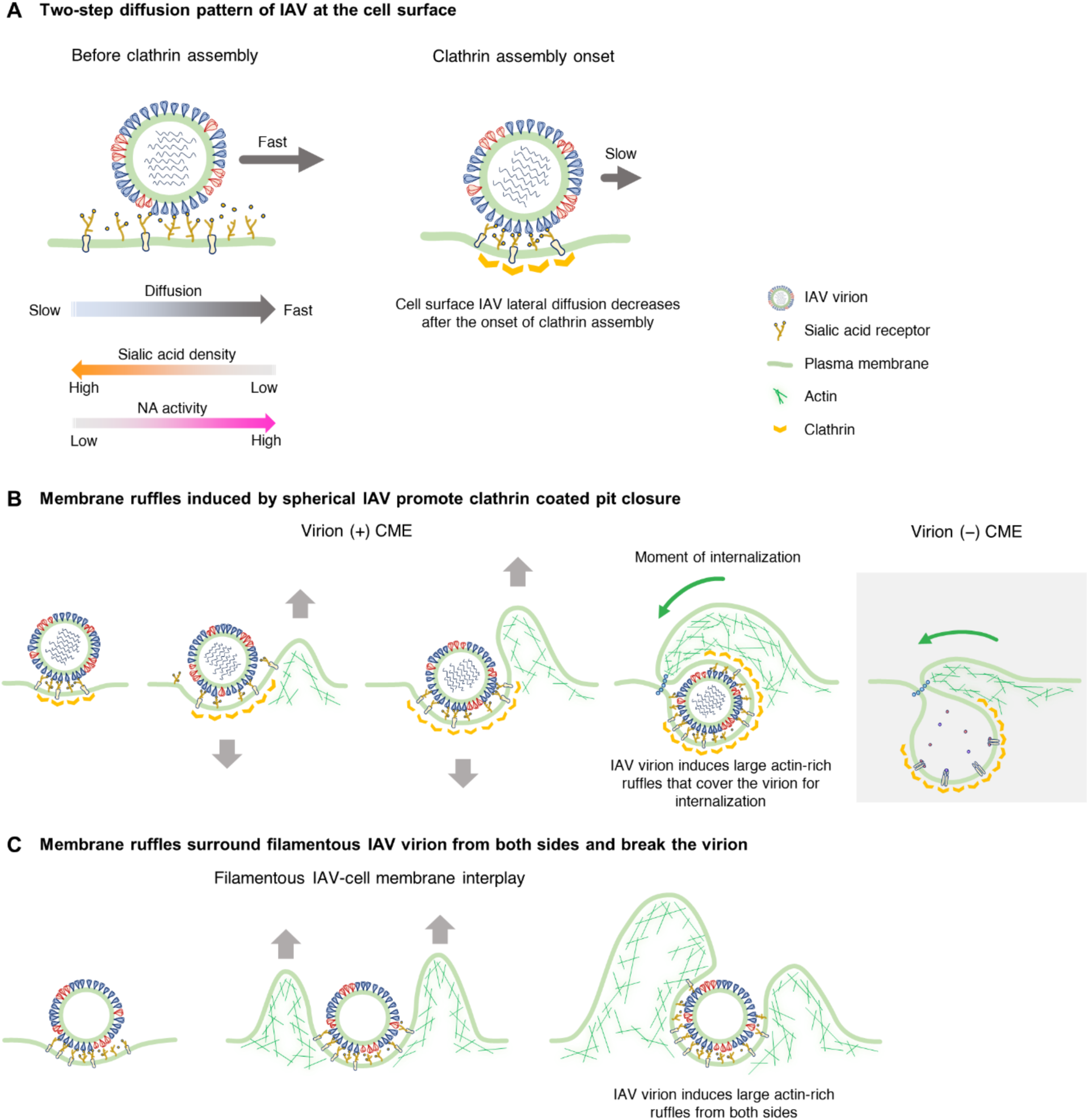
Model of IAV diffusion and membrane interaction leading up to clathrin-mediated endocytosis. **(A)** Two-step diffusion pattern of IAV at the cell surface. In the absence of clathrin assembly (left), IAV lateral diffusion depends on sialic acid density and neuraminidase activity. The lower the sialic acid density, the larger the IAV diffusion. The lower the neuraminidase activity, the smaller the IAV diffusion. Upon the onset of clathrin assembly (right), the subsequent IAV diffusion decreases. **(B)** Morphological changes of plasma membrane during spherical IAV CME. An actin-rich membrane ruffle is formed following clathrin assembly and pulls the membrane up and the virion down. Eventually, the virion is enveloped by membrane ruffles that are larger than those in virion (-) CME. **(C)** Interplay between filamentous IAV and the live-cell membrane. An actin-rich membrane ruffle is formed around the middle of the filament and surround the virion from both sides, leading to filament fragmentation.

**Movie S1. Morphological changes of the MDCK cell surface.**

A representative movie of the cell membrane morphological changes captured using AFM, related to Fig. 1B. Sequential images were acquired every 10 s at 27°C. The AFM image is a topographic image, with the height direction represented by 400 nm: bright areas are higher and dark areas are lower. Image size on the x-y plane: 6.0 × 4.5 μm^2^. Membrane ruffles of various sizes were confirmed, moving dynamically on the cell surface.

**Movie S2. IAV lateral diffusion on MDCK cell surface.**

A representative movie of Alexa Fluor 594-labeled virions diffusing on the cell membrane captured using the ViViD-AFM, related to Fig. 1D. Sequential images were acquired every 10 s at 27°C. Movies are shown in the following order: fluorescence movie of IAV-AF594 by confocal microscopy, morphology movie of virion by AFM, and integrated movie of fluorescence and morphology. Image size on the x-y plane: 6.0 × 4.5 μm^2^. The height range of AFM: 400 nm. Two virions diffusing on the cell membrane were indicated by arrowheads.

**Movie S3. IAV diffusion on MDCK cell surface before clathrin assembly.**

A representative morphological movie (from t=225 s to t=575 s) of the virion diffusion on the cell membrane before the clathrin assembly captured using ViViD-AFM, related to Fig. 2C-E. Sequential images were acquired every 5 s at 37°C. The tracking virion was indicated by arrowhead. AFM image size on the x-y plane: 6.0 × 4.5 μm^2^. The height range of AFM: 400 nm.

**Movie S4. IAV diffusion on MDCK cell surface before and after clathrin assembly.**

The movie (from t=0 s to t=80 s) shows virion diffusion on the cell membrane before and after the onset of clathrin assembly (t=50 s) captured using the ViViD-AFM, related to Fig. 3A-D. Sequential images of morphology and fluorescence of Alexa Fluor 594-labeled virion and CLCa -EGFP were acquired every 5 s at 37°C. In the movie, morphology image of the virion and cell membrane (left), the fluorescence image of IAV-AF594 (magenta) (middle) and CLCa -EGFP (green) (right) were displayed. Arrowheads indicate the position of the virion in the movie of AFM and the position of IAV-AF594 in the movies of IAV-AF594, and the position of clathrin assembly in the movie of CLCa-EGFP, respectively. Image size on the x-y plane: 6.0 × 4.5 μm^2^. The height range of AFM: 400 nm. At the arrowhead position in the right fluorescence image, clathrin assembly was observed from t=50 s.

**Movie S5. Actin-rich membrane ruffles promote IAV internalization into MDCK cell.**

A representative movie of Alexa Fluor 594-labeled virion internalization into the cell, accompanied by EGFP-Lifeact-positive membrane ruffles captured using the ViViD-AFM, related to Fig. S13. Sequential images of morphology and fluorescence were acquired every 10 s at 27°C. In the movie, morphology image of the virion and cell membrane (left), merged fluorescence image of EGFP-Lifeact (green) and IAV-AF594 (magenta) (middle), and merged image of AFM and fluorescence (right) were displayed. The virion on the cell membrane was indicated by arrowheads. Moment of IAV internalization (t = 0 s) was set as the time when the virion disappeared from AFM images but was detectable by fluorescence. Image size on the x-y plane: 1500 × 1500 nm^2^. The height range of AFM: 400 nm.

**Movie S6. Moment of IAV internalization into MDCK cell.**

A representative morphological 3D movie of virion internalization captured using ViViD-AFM, related to Fig. S14. Sequential images were acquired every 10 s at 27°C. The virion was colored magenta. At t = 0 s, the virion was covered by a membrane ruffle and internalized into the cell.

**Movie S7. CME of IAV preceded by multiple cycles of clathrin assembly.**

The movie (from t=-585 s to t=45 s) of the virion-membrane interplay accompanied with clathrin assembly multiple times captured using ViViD-AFM, related to Fig. S14. Sequential images were acquired at 37°C every 5 s. The virion disappeared from the cell surface at t = 0 s. The tracking virion was colored magenta. AFM image size on the x-y plane: 6.0 × 4.5 μm^2^. The height range of AFM: 400 nm.

**Movie S8. Tracking of IAV virion before and after CME into MDCK cell.**

The movie (from t=-230 s to t=180 s) of the virion trajectory before and after the virion internalization (t=0 s) into the cell captured using ViViD-AFM, related to Fig. S14. Sequential images of morphology and fluorescence of CLCa -EGFP and Alexa Fluor 594-labeled virion were acquired at 37°C every 5 s. In the movie, the merged fluorescence image of IAV-AF594 (magenta) and CLCa -EGFP (green) (left), morphology image of the virion and cell membrane (middle) and merged image of fluorescence and AFM (right) were displayed. The virion morphology visible in the AFM image is indicated by an arrowhead. In fluorescence images, the trajectory of the virion indicated with arrowhead is overlaid, and trajectories in extracellular and intracellular environments are colored orange and gray, respectively. Fluorescence image (left) size on the x-y plane: 6.0 × 8.6 μm^2^. AFM image (middle) and merged image (right) size on the x-y plane: 6.0 × 4.5 μm^2^. The height range of AFM: 400 nm.

**Movie S9. Morphological deformation of filamentous IAV virions by membrane ruffles.**

The movie (from t=0 s to t=160 s) of filamentous virion-cell membrane interplay captured by ViViD-AFM, related to Fig. 5. Sequential images of morphology and fluorescence of DiI were acquired at 27°C every 5 s. In the movie, morphology image of virion and cell membrane (left), fluorescence image of IAV-DiI (middle), and the merged image of AFM and fluorescence (right) are displayed. Scale bar: 500 nm. The height range of AFM: 400 nm.

**Movie S10. Fragmentation of filamentous IAV followed by internalization and intracellular transport.**

The movie (from t=0 s to t=1070 s) of filamentous virion in MDCK cells captured by confocal microscope, related to Fig. S16. Sequential fluorescence images of IAV-DiI were acquired at 27°C every 10 s and were displayed as a movie. Arrowheads indicate the tip of the filament separating and moving away from the main filament. Scale bar: 5 µm.

